# Pairwise Interactions in Adjuvant Combinations Dictate Immune Responses and Inform Cancer Immunotherapy Design

**DOI:** 10.1101/2020.07.11.198879

**Authors:** Surya Pandey, Adam Gruenbaum, Tamara Kanashova, Philipp Mertins, Philippe Cluzel, Nicolas Chevrier

**Affiliations:** Pritzker School of Molecular Engineering, The University of Chicago, Chicago, IL 60637, USA; Proteomics Platform, Max Delbrück Center for Molecular Medicine in the Helmholtz Society, and Berlin Institute of Health, 13125 Berlin, Germany; School of Engineering and Applied Science & Department of Molecular and Cellular Biology, Harvard University, Cambridge, MA 02138, USA

## Abstract

The immune system makes decisions in response to complex combinations of microbial inputs. We do not understand the combinatorial logic that governs how the interplay between higher-order combinations of microbial or adjuvant signals shape immune responses, which hampers the rational design of vaccines and immunotherapies. Here, using *in vitro* coculture experiments and statistical analyses, we discover a general property for the combinatorial sensing of microbial signals, whereby the effects of triplet combinations of adjuvants on immune responses can be explained by the effects of single and pairwise stimulations. Mechanistically, we find that adjuvant singles and pairs dictate the information signaled by triplets in mouse and human DCs at the levels of transcription, chromatin and protein secretion. Furthermore, we exploit this simplifying property to develop and characterize cell-based immunotherapies using adjuvant combinations with anti-tumor properties in mouse models. We conclude that the processing of complex mixtures of microbial or adjuvant inputs by immune cells is governed by pairwise effects, which will inform the rationale combination of immunomodulatory agents such as adjuvants to manipulate immunity.

## INTRODUCTION

Biological systems make decisions in response to complex combinations of signals. For example, to stop an infection, the immune system has learned to recognize and exploit the inter-dependencies of microbial signals by evolving in a chance-driven world of encounters with pathogens. By mimicking such responses to complex microbial signals, live vaccines that are empirically attenuated from pathogens have been a powerful means to yield life-long immunity against many deadly pathogens (Plotkin et al., 2017). However, the rational design of non-live vaccines using immunomodulatory agents such as adjuvants has remained an elusive task in many cases where live vaccination is not efficacious or feasible (Coffman et al., 2010; Levitz and Golenbock, 2012; Pulendran and Ahmed, 2011). To tackle this challenge, a central question to answer is how do complex combinations of microbial or adjuvant signals shape immune responses? Filling this fundamental gap in our knowledge is critical to learn how to rationally choose and combine adjuvants to manipulate immunity against infectious and non-infectious diseases such as cancer.

The molecular signals derived from pathogens, live vaccines or adjuvants are largely processed by pattern recognition receptors (PRRs) of the innate immune system (Ablasser and Chen, 2019; Brown et al., 2018; Chow et al., 2018; Iwasaki and Medzhitov, 2015; Janeway, 1989; Medzhitov, 2009; Takeuchi and Akira, 2010). To date, pathogen-sensing pathways have been much studied one pathway at a time. As a first step beyond the analysis of single pathway effects, many examples of synergy, independence or antagonism between pathogen-sensing pathways have been reported using pairwise stimulations, adjuvant combinations or genetic deletions (Bagchi et al., 2007; Cappuccio et al., 2015; Crozat et al., 2009; Elinav et al., 2011; Gantner et al., 2003; Kasturi et al., 2011; Kawai and Akira, 2011; Lin et al., 2017; Loo and Gale, 2011; Napolitani et al., 2005; Negishi et al., 2012; Nish and Medzhitov, 2011; Osorio and Reis e Sousa, 2011; Ozinsky et al., 2000; Thaiss et al., 2016). These observations suggest that responses to two microbial stimuli cannot be explained by combining the effects of single stimuli. In addition, the higher-order effects of microbial inputs on pathogen-sensing pathways and downstream immune responses have not been systematically analyzed. Thus, we do not know how pathogen-sensing pathways respond collectively to complex mixtures of input signals, as is the case with natural infections or mixtures of adjuvants in vaccines.

Since studying the full scope of combinatorial effects in pathogen sensing is impractical because the number of experiments grows exponentially with the number of stimuli, we need an innovative strategy to decipher the complexity and the combinatorial logic underlying innate immune sensing. The key challenges to address include (1) identifying assays and readouts to capture the complexity of multi-input effects on the immune response both *in vitro* and *in vivo* for immunotherapeutic design; (2) dissecting the mechanistic underpinnings of combinatorial sensing; and (3) finding ways to predict higher-order effects in cells and in the host, as a means to circumvent the need for testing many combinations.

Here, we studied how the interplay between higher-order combinations of microbial input signals shapes the output of immune responses. Specifically, we asked if the relationships between the effects of singles, pairs, and triplets of inputs can reveal the combinatorial logic governing microbial sensing by the immune system (**Figure 1A**). First, we measured the effects of a representative set of seven microbial stimuli and all corresponding pairwise (21) and triplet (35) combinations on T cell responses using dendritic cell (DC)-T cocultures *in vitro*. We reasoned that our DC-T coculture system provides an integrated readout for the various DC-derived signals that are regulated by the combinatorial activation of pathogen-sensing pathways. Remarkably, we found that the effects of triplet combinations of stimuli on DC-T responses can be explained by the effects of single and pairwise stimulations. Second, as a mechanistic basis for our finding, we observed that singles and pairs of microbial inputs dictate the information signaled by triplets in mouse and human DCs at the levels of transcription, chromatin and protein secretion. Third, we asked if the combinatorial logic governing pathogen sensing *in vitro* would be applicable *in vivo* using cell-based immunotherapies in mouse models of cancer. We identified several triplets of immune adjuvants with potent anti-tumor effects and showed that their effects can be explained using the *in vivo* effects of adjuvant singles and pairs only. Overall, we discovered a general property that simplifies the combinatorial logic of pathogen sensing and can be exploited to rationally combine adjuvants for therapeutic design.

**Figure 1.**
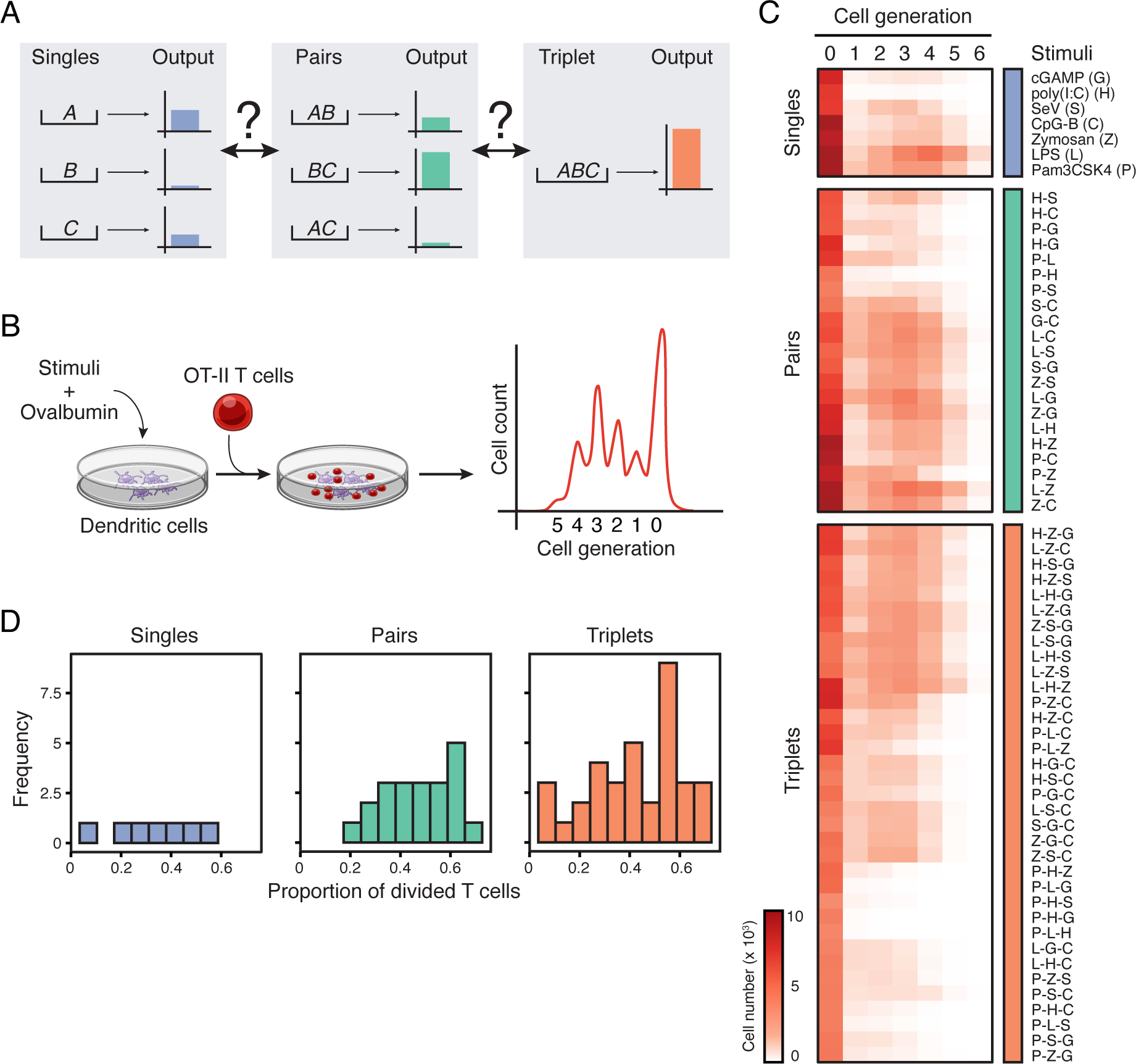
Systematic analysis of the combinatorial effects of microbial stimuli in a coculture system. (A) Schematic depicting the unknown relationships between the effects of input singles (*A*, *B* and*C*; left), pairs (*AB*, *AC* and *BC*; center) and triplet (*ABC*; right) on a hypothetical output response. (B) Schematic overview of the coculture assay used to measure the combinatorial effects of microbial inputs. Dendritic cells stimulated with singles, pairs or triplets of stimuli are pulsed with ovalbumin and subsequently incubated with transgenic OT-II T cells. T cell proliferation is measured by CFSE dilution. (C) Heatmap showing all 63 microbial stimuli combinations (rows) used in the study: 7 singles (blue; top), 21 pairs (green; middle) and 35 triplets (orange; bottom). Columns indicate the cell generation and values are average cell numbers (color scale) (*n* = 8). (D) Frequency distribution of the proportion of divided T cells for single (left), pair (center) and triplet (right) ligand stimulations.

## RESULTS

### A coculture assay to characterize the combinatorial effects of microbial stimuli

First, we sought to explore the combinatorial effects of microbial inputs on the output of an immune response (**Figure 1A**). To this end, we systematically measured the effects of singles, pairs and triplets of microbial stimuli on the ability of DCs to instruct T cell responses (**Figure 1B**). We reasoned that measuring T cell proliferation provides an integrated readout for the various DC-derived signals triggered by the combinatorial activation of pathogen-sensing pathways, which include the regulation of antigen presentation and membrane-bound and secreted co-stimuli such as cytokines.

We selected seven well-established ligands for pathogen-sensing pathways: (*i*) lipopolysaccharide (LPS or L), (*ii*) Pam3CSK4 (P), (*iii*) high molecular weight poly(I:C) (H), (*iv*) CpG DNA type B (C), (*v*) Sendai virus (SeV or S), (*vi*) depleted zymosan (Z), and (*vii*) cyclic [G(3’,5’)pA(3’,5’)p] (3’3’-cGAMP or G), which are agonists for, respectively, TLR4, TLR2, TLR3/MDA-5, TLR9, RIG-I, Dectin-1 and STING, the adaptor downstream of the cGAS pathway. These ligands were selected (1) to encompass most PRR families, (2) because their receptors are expressed in steady-state DCs (**Figure S1A**) and functional as shown using knockout cells (**Figure S1B**), (3) by performing dose-response experiments (**Figure S1C**), and (4) ensuring that each ligand had no direct effects on T cell viability and proliferation (**Figure S1D**). Next, we incubated mouse bone-marrow-derived DCs with the chicken ovalbumin (OVA) protein with or without ligands for six hours, then washed and cultured DCs with carboxyfluorescein succinimidyl ester (CFSE)-labeled OT-II transgenic CD4^+^ T cells specific for the OVA_323-339_ peptide bound to the major histocompatibility (MHC) class II molecule I-A^b^ (**Figure 1B**) (Barnden et al., 1998). In these experimental settings, ligands P and L were the strongest inducers of T cell proliferation (>45%), G and H the weakest (<10%), and C, S and Z showed intermediate levels (20-40%) (**Figure S2A**). We then measured the effects of all possible two- and three-way ligand combinations in our DC-T coculture system, leading to 21 pairs and 35 triplets in total (**Figure 1C** and **Figure S2A** and **Table S1**). The distributions of T cell proliferation values across singles, pairs and triplets were comparable, albeit slightly shifted towards higher values (>60% of T cells divided) in pairs and triplets (**Figure 1D** and **Figure S2B**). We also found that IFN-γ secretion by T cells was higher in pairs compared to singlets, and seemingly reached a plateau at the pair and triplet levels (**Figure S2C**). Importantly, these changes in proliferation and IFN-γ were not due to toxicity effects of ligand combinations: 90% (57/63) of the conditions tested led to >65% viability in T cells, with the exception of 6 triplets, namely P-Z-G, P-Z-S, P-H-Z, P-L-Z, Z-S-C and Z-G-C, which led to viability values ranging from 39 to 62% (**Figure S2D**).

### Pairwise and single stimuli effects suffice to accurately describe triplet effects

Next, we examined the relationship between single and pairwise ligand stimulations and the net interactions of three ligands using the T cell growth data from our combinatorial experiments (**Figure 1C** and **Table S1**). First, we sought to classify how pairs of microbial signals interact. Qualitatively, pairwise ligand stimulations of DCs resulted in T cell growth patterns which appeared synergistic, antagonistic or additive as shown, for example, in the pairs of ligands that compose the following triplets: Z-S-G, P-S-G and P-L-Z (**Figure 2A**). We ensured that pairwise ligand effects were due to the integration of signals at the level of single DCs, as shown by experiments mixing (1) ligands to stimulate DCs, or (2) DCs stimulated with single ligands (**Figure S3A**). Indeed, mixing cells stimulated with single ligands did not lead to the synergistic effects observed when mixing ligands for 10 out of the 15 pairs of ligands tested (**Figure S3B**). To quantify the effects of crosstalk between ligand pairs, we used the proliferation index (*Pi*) as a proxy for the T cell response in our *in vitro* coculture system, which corresponds to the average number of divisions per activated T cell in our DC-T coculture system (Roederer, 2011) (**STAR Methods**). Using the proliferation indices associated with two ligands, *A* and *B*, used alone, *Pi_A_* and *Pi_B_*, and in combination, *Pi_AB_*, we computed a pairwise interaction score *I_AB_* for each ligand pair defined as *I_AB_* = *Pi_AB_* – *Pi_A_Pi_B_* (**STAR Methods**). The pairwise interaction score *I_AB_* quantifies the level of synergy, antagonism or lack thereof that results from crosstalk between two pathways triggered by the ligands *A* and *B*. Indeed, if two responses are independent as postulated for example in the Bliss independence model (Bliss, 1939; Bliss, 1956), then *I_AB_* would be equal to *Pi_A_Pi_B_* and the effects of ligands *A* and *B* would simply be deemed independent from one another. However, any deviations of *I_AB_* from *Pi_A_Pi_B_* indicates the existence of interactions between the ligand-activated pathways, which can either enhance (synergy) or inhibit (antagonism) each other’s effects. For example, the pairs P-L and P-S showed antagonistic effects whereas Z-S, Z-C and S-G were synergistic (**Figure 2B**). Overall, pairwise interaction score calculations revealed that 52% (11/21) of the pairs were synergistic, 10% (2/21) were antagonistic and 38% (8/21) were additive (**Figure 2D**), which indicates a complex interplay between pairs of ligands at the level of DC-T cell responses.

**Figure 2.**
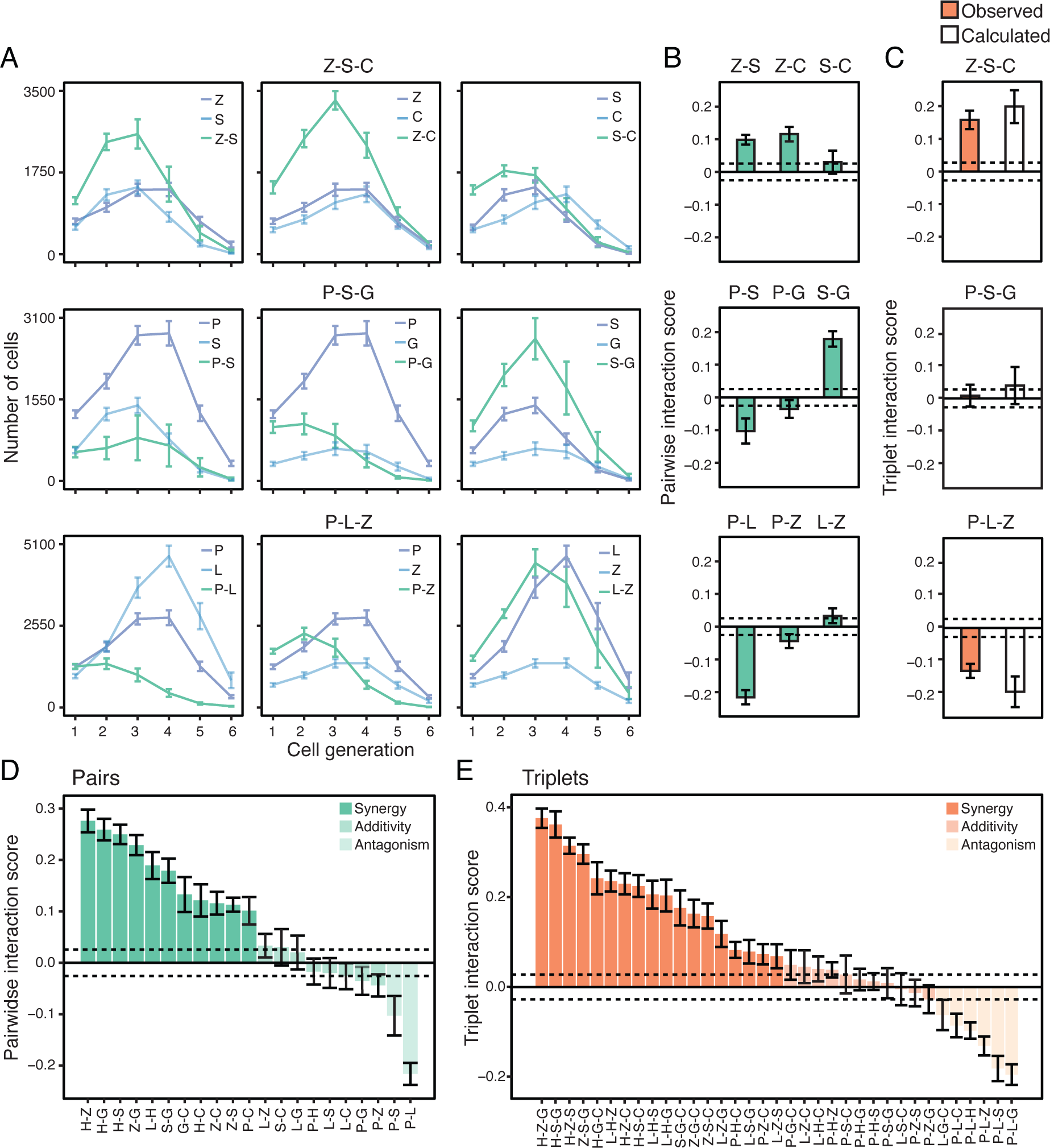
Pairwise effects qualitatively combine to explain the net effect of triplets of microbial stimuli. (A) T cell growth pattern upon DC stimulations with ligand singles and pairs. Line plots showing the number of OT-II cells in each generation of activated cells (cell generation 1 to 6) upon coculture with DCs stimulated with singles (blue) and pairs (green) corresponding to the triplets Z-S-C (Zymosan/SeV/CpG-B; top), P-S-G (Pam3CSK4/SeV/cGAMP; middle) and P-L-Z (Pam3CSK4/LPS/Zymosan; bottom). Error bars, SEM (*n* = 8). (B-C) Pairwise (*I_AB_* = *Pi_AB_* - *Pi_A_Pi_B_*; b) and triplet (*I_ABC_* = *Pi_ABC_* - *Pi_A_Pi_B_Pi_C_*; c) interaction scores for the triplets Z-S-C (top), P-S-G (middle) and P-L-Z (bottom) (C) and their composite pairs (B). Triplet scores (C) are derived from *Pi_ABC_* values observed experimentally (orange) or calculated (white) using an Isserlis formula (*Pi_ABC_* = *Pi_AB_Pi_C_+ Pi_AC_Pi_B_* + *Pi_BC_Pi_A_* - 2*Pi_A_Pi_B_Pi_C_*). Dashed horizontal lines indicate the mean of SEM values across all pairs (B) or triplets (C), used as thresholds for synergy and antagonism (+/-0.0256 for pairs and +/-0.0275 for triplets). Error bars, SEM (*n* = 8). (D-E) Bar plots showing pairwise (*I_ab_* = *Pi_ab_*-*Pi_a_Pi_b_*; D) and triplet (*I_abc_* = *Pi_abc_*-*Pi_a_Pi_b_Pi_c_*; E) interaction scores for indicated ligand combinations. Dashed horizontal lines indicate the mean of SEM values across all pairs (D) or triplets (E) and are used as thresholds for synergy (opaque colors), additivity (medium opacity colors) or antagonism (light opacity colors). Error bars, SEM (*n* = 8).

Second, we investigated the relationship between the net effects of three-ligand interactions and matching singles and pairs. To do so, we computed a triplet interaction score *I_ABC_* for each triplet of ligands defined as *I_ABC_* = *Pi_ABC_* – *Pi_A_Pi_B_Pi_C_* (**STAR Methods**), which encapsulates the level of net pairwise and triplet interactions by subtracting single ligand effects from the triplet proliferation index. The proportions of synergistic (51%, 18/35), antagonistic (17%, 6/35) and additive (31%, 11/35) triplet effects were comparable to those measured for pairwise effects (**Figure 2E**). Remarkably, pairwise effects combined qualitatively in a variety of ways to yield triplet effects *I_ABC_*. For example, synergistic interactions of intermediate strength between pairs, such as those between Z-S and Z-C, can combine to yield a cumulative effect that is strongly synergistic (**Figure 2B-C**, top panel). Conversely, pairs with antagonistic (P-S and P-G) and synergistic (S-G) interactions can combine to yield a cumulative triplet effect (P-S-G) that is close to zero (**Figure 2B-C**, middle panel). Further, the pairs P-Z and L-Z showing weak interactions in negative and positive directions, respectively, combined with the strongly antagonistic pair P-L to generate a net P-L-Z triplet effect whose magnitude is similarly antagonistic than that of the P-L pair alone (**Figure 2B-C**, bottom panel). Taken together, these results qualitatively showed that pairwise effects combine in a variety of ways to explain the net effect of triplets, suggesting an intrinsic property for pathogen-sensing pathways whereby three-way interactions between input signals may be encapsulated in the corresponding pairwise effects when monitoring DC-T coculture outputs.

To test the robustness of this property, we asked whether triplet ligand interactions could be predicted by using only data from single and pairwise effects. To do so, we used a statistical analysis derived from the Isserlis theorem (Isserlis, 1918), which was previously applied to evaluate higher-order effects in combinations of antibiotics on bacterial growth (Wood et al., 2012). We used the following Isserlis expression: *Pi_ABC_* = *Pi_AB_Pi_C_* + *Pi_AC_Pi_B_* + *Pi_BC_Pi_A_* − 2*Pi_A_Pi_B_Pi_C_*, where *Pi* is the proliferation index of T cells from DC-T cocultures for the indicated combinations of ligands *A*, *B* and *C* (**STAR Methods**). In this approach, the equality is satisfied when there is no three-way interactions between signals. As a first approximation, we compared the triplet interaction scores obtained from experimental values to the scores inferred computationally using the Isserlis statistical approach and found a strong agreement between observed and calculated values (R^2^ = 0.9) (**Figure 2C and 3A**). Going further, we found that the proliferation indices for triplets obtained from experiments were strikingly similar to those calculated using the Isserlis formula that uses only single and pairwise proliferation indices as input values (R^2^ = 0.77) (**Figure 3B**). The statistical significance of this correlation between values observed experimentally and calculated with Isserlis was confirmed by bootstrap analysis from one million trials of randomly scrambling singles and pairs across triplets of different ligands (**Figure 3C** and **STAR Methods**). Importantly, using an additional transgenic CD4^+^ T cell model specific for an H2-I-A^b^-restricted lymphocytic choriomeningitis virus (LCMV) glycoprotein-derived epitope (residues 61 to 80) (Oxenius et al., 1998), we obtained similar results suggesting the broad applicability of our observation (**Figure 3A-C**, bottom panels).

**Figure 3.**
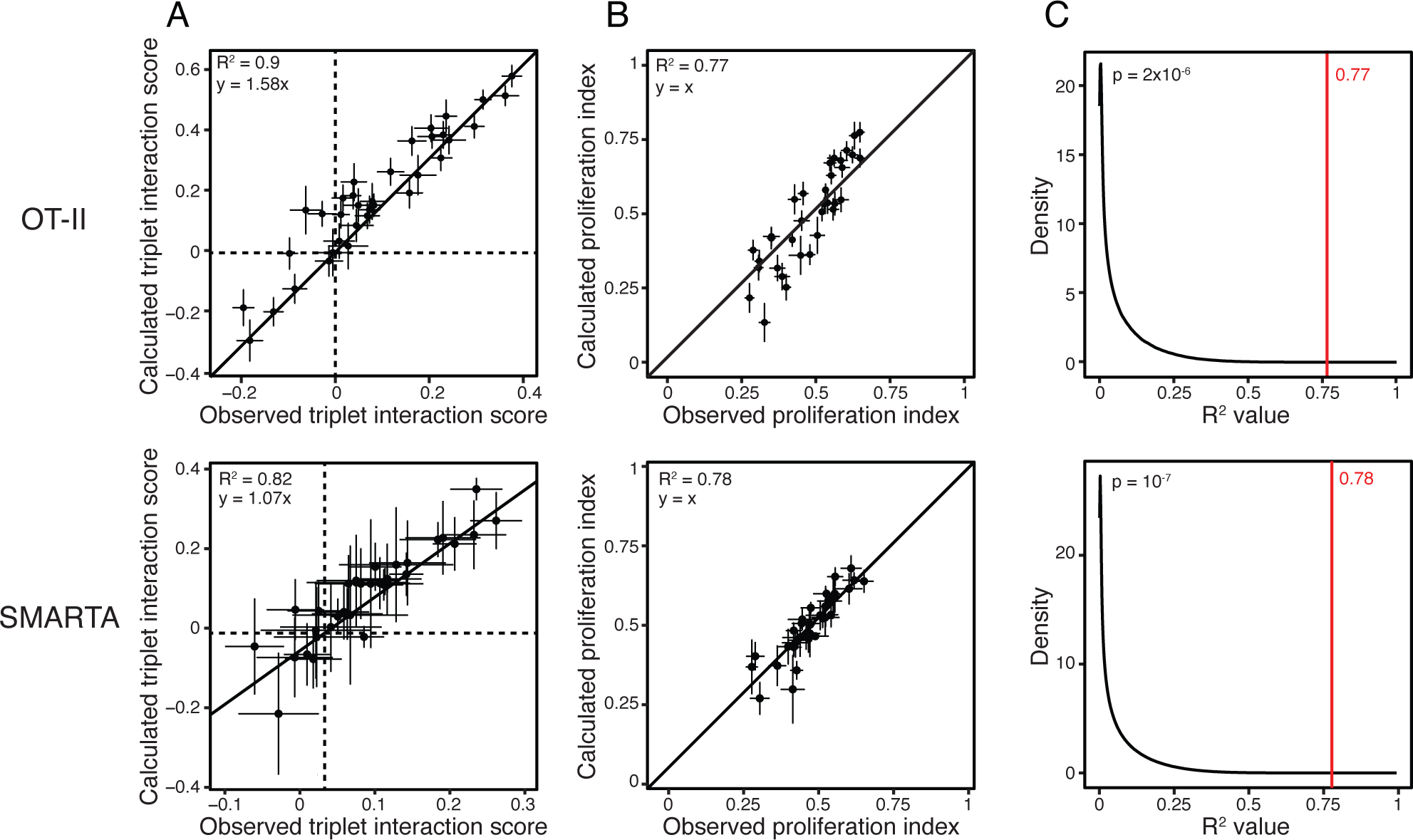
Single and pairwise stimulations predict higher-order effects. (A) Dot plots of the observed (X axis) and calculated (Y axis) triplet interaction scores for all 35 ligand triplets tested and using CD4^+^ T cells from OT-II (top) or SMARTA (bottom) transgenic mice. Dashed lines indicate the X and Y axes. The solid line indicates a linear model fit (top, y = 1.58x; bottom, y = 1.07x). Error bars, SEM (*n* = 8 for OT-II and *n* = 2 for SMARTA). (B) Dot plots of the observed (X axis) and calculated (based on the Isserlis formula; Y axis) triplet proliferation index values for all 35 ligand triplets tested, using OT-II (top) or SMARTA (bottom) T cells. The solid line indicates y = x. Error bars, SEM (*n* = 8 for OT-II and *n* = 2 for SMARTA). (C) Distribution of R^2^ values obtained between observed and calculated proliferation indices for ligand triplets with random shuffling by bootstrapping (black line) or without (red line). The number in red indicates the R^2^ value obtained by correlating experimental and calculated triplet proliferation indices as shown in B. Data obtained with OT-II (top) and SMARTA (bottom) T cells.

Using indices that capture other characteristics of T cell growth patterns from CFSE profiles led to poor correlations between observed and calculated values, with the exception of the replication index and, to a lesser extent, the expansion index which capture similar information to *Pi* (**Figure S3C** and **STAR Methods**) (Roederer, 2011). Using IFN-γ production was not a good predictor of triplet effects using only data from singles and pairs (**Figure S3D**). In addition, the types of PRR pathways targeted in ligand triplets did not impact the distribution of proliferation indices, although combining ligands from two or three different PRR families of receptors tended to display higher proliferation level than those combinations of ligands targeting only the Toll-like receptor (TLR) family (**Figure S3E-F**).

Furthermore, we showed in two control experiments that the heterogeneity of bone marrow-derived DC cultures previously described by others (Helft et al., 2015), did not impact our findings about estimating triplet effects using only data from singles and pairs. First, the three cell subsets present in DC cultures based on the surface markers CD11b, CD11c and MHC class II (**Figure S3G**), responded similarly to combinatorial stimulations as measured by the transcriptional expression of four signature cytokines of DC activation (*i.e.*, *Cxcl1*, *Cxcl10*, *Ifnb1*, *Il6*), albeit with variations in the magnitude of gene activation (**Figure S3H**). Second, when using a homogenous DC culture based on the enrichment of MHC class II-positive cells (**Figure S3I**), we found that the proliferation indices for triplets obtained experimentally were strikingly similar to those calculated using the Isserlis formula (**Figure S3J**).

Overall, these results reveal a simplifying property for the combinatorial sensing of microbial signals, whereby the effects of triplet combinations of stimuli can be reduced to the effects of single and pairwise stimulations.

### Singles and pairs of microbial stimuli dictate the information signaled by triplets in DCs

To investigate the effects of combinatorial stimulations on DC states, we first measured changes in gene expression in DCs stimulated for 6 hours with all the ligand singles (7), pairs (21) and triplets (35) used in our co-culture assay. In total, we identified 1,357 genes differentially expressed between stimulated and control, unstimulated DCs across all 63 combinations (**Figure 4A, Table S2A, STAR Methods**). Interestingly, all of the transcriptional profiles for triplets clustered closely with singles and pairs using hierarchical clustering (**Figure 4A**) and principal component analysis (PCA) (**Figure 4B**), which suggests that the transcriptional states of DCs stimulated with ligand triplets is comparable to that of DCs exposed to singles and pairs. To test this, we counted how many genes were found to be up- or down-regulated in a ligand pair but not in either one of the singles forming that pair (**STAR Methods**). We found that 67% (14/21) of the pairs tested regulated genes that were not differentially expressed in the corresponding single-ligand treated cells compared to unstimulated control cells (**Figure 4C, Figure S4A, Table S2B**). 0.2 to 39% of the genes regulated by those 14 pairs were found to be newly regulated at the pair level (**Figure 4C**). In contrast, 77% (27/35) of the triplets tested did not regulate new genes relative to matching singles and pairs, with only 20% (7/35) of the triplets displaying 0.2-1.3% of newly regulated genes and one triplet, P-H-Z, regulating 26 new genes, which is 6.3% (26/412) of all its regulated genes (**Figure 4C, Figure S4A-I, Table S2B**).

**Figure 4.**
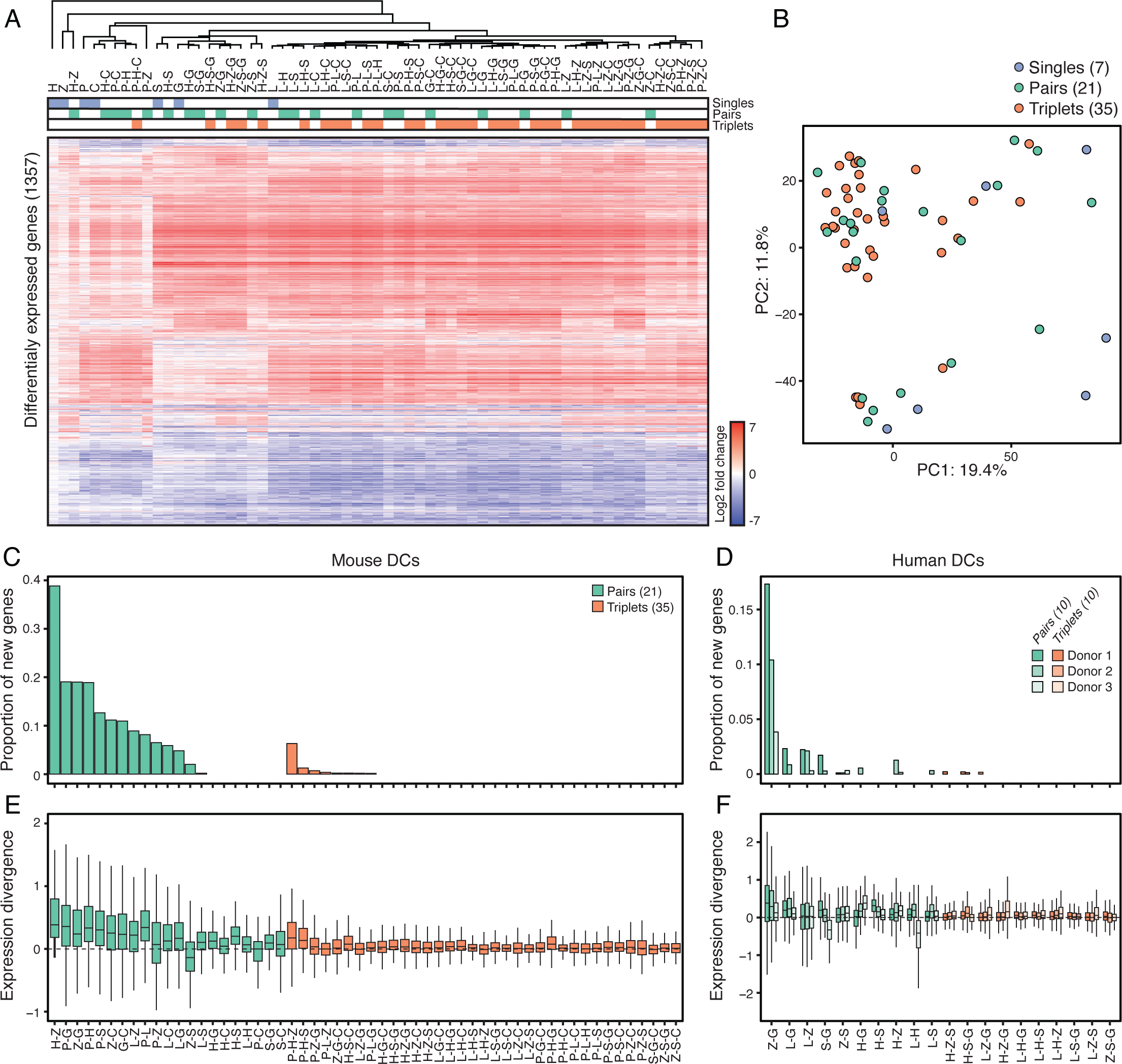
Single and pairwise stimulations dictate the information signaled by triplets in mouse and human DCs. (A) Heatmap of differentially expressed genes (1,357 in total; rows) from mRNA profiles of mouse DCs stimulated with ligand singles (blue), pairs (green) or triplets (orange) (columns). Values are log2 fold-changes relative to unstimulated cells (FDR-adjusted p value < 0.01; *n* = 3). Hierarchical tree is based clustering using Pearson’s correlation. (B) Principal component analysis (PCA) of mRNA profiles of mouse DCs stimulated with ligand singles (7; blue), pairs (21; green) or triplets (35; orange). (C-D) Bar plots showing for each ligand pair (green) or triplet (orange) the proportion of genes regulated by a triplet or a pair but not by matching composite singles and/or pairs relative to the total number of genes regulated by that triplet, using mouse (C) and human (D) DCs (*n* = 3). (E-F) Box plots showing the smallest divergences between the level of expression of genes regulated by ligand pairs (green) and triplets (orange) and their composite singles and/or pairs, using mouse (E) and human (F) DCs (*n* = 3). Box plots represent the median, first quartile and third quartile with lines extending to the furthest value within 1.5 of the interquartile range (IQR).

To test whether the genes that were regulated by triplets showed changes in expression levels that were similar or different in amplitude compared to all matching singles and pairs, we computed the smallest possible difference in fold-change values relative to control cells for a given gene between (1) a pair and its matching singles, and (2) a triplet and its matching singles and pairs, which we refer to as expression divergence (**STAR Methods**). A low expression divergence value indicates that the amplitude of the expression change for a given gene upon triplet stimulation is close to at least one of the composite stimuli for that triplet, whereas a high expression divergence highlights genes whose change in expression upon triplet stimulation is different from all composite stimuli singles and pairs. While pairs led to significant changes in expression levels compared to their matching singles, triplets triggered little to no change in expression levels compared to their composite single and pairwise conditions (**Figure 4E**). These results show that the genes regulated by ligand triplets are also found to be regulated in matching singles and pairs and at similar levels relative to unstimulated cells. In other words, triplets of ligands do not seem to encode new information at the transcriptional level compared to singles and pairs. Importantly, similar results were obtained when using DCs derived from human blood monocytes isolated from three independent donors, suggesting that this property of the response of PRR pathways to multi-stimuli stimulations is preserved in humans (**Figure 4D and 4F, Figure S5A, Table S3**).

Next, we tested whether this phenomenon would hold true at the chromatin and secreted protein levels. To do so, we first compared changes in the gene expression and genome-wide chromatin accessibility states of DCs stimulated with three randomly selected ligand triplets P-L-H, P-S-G or Z-S-G and their matching singles and pairs (**STAR Methods**). We found that, similarly to what we observed at the mRNA level, triplet stimulations triggered changes in chromatin accessibility at genomic loci that were already regulated by single and pairwise stimulations (**Figure S5B-C, E-F and H-I, Table S4**). Second, we used mass spectrometry on cell culture supernatants to measure changes in the secretome of DCs stimulated with the same three triplets and their matching singles and pairs (**Figure S5D and G, Table S5, STAR Methods**). Similar to the transcriptional and chromatin levels, the secretome of DCs did not differ significantly in triplet conditions compared to matching singles and pairs (**Figure S5B-G**). Taken together, these observations suggest a model whereby activating triplets of pathogen-sensing pathways does not lead to the regulation of new genes – at the level of chromatin, transcription and protein secretion – compared to corresponding singles and pairs.

### Adjuvant triplets generate potent DC-based vaccines against melanoma in mice

Having identified a simplifying, intrinsic property that explains the collective effects of pathogen-sensing pathways in cocultures *in vitro*, we next sought to test the applicability of this property in the natural setting of the host. To do so, we sought to establish an *in vivo* model in which triplets of PRR ligands would lead to a variety of outcomes in terms of host protection. We reasoned that finding triplets leading to different outcomes for the host – such as protection against tumor versus none – would allow us to test whether using data from single and pairwise adjuvant treatments could describe triplet effects *in vivo*. To test this, we used a DC vaccination model to precisely control the exposure of DCs to combinations of microbial stimuli prior to injecting DCs subcutaneously in mice to assess their protective potential. We selected 12 out of the 35 triplets studied to cover all major transcriptional clusters observed *in vitro* (**Figure 4A**). We generated DC vaccines by stimulating DCs *ex vivo* with either one of the triplets for 6 hours in the presence of the full ovalbumin (OVA) protein in the culture medium. Next, we screened the effects of these DC-based vaccines by injecting them subcutaneously in mice which received 10^5^ OVA-expressing B16.F10 melanoma cells (B16-OVA) in the contralateral skin region (**Figure 5A**). Remarkably, out of the 12 triplets used to create DC vaccines, 4 led to a strong decrease in melanoma growth whereas the remaining 8 triplets did not (**Figure 5B**).

**Figure 5.**
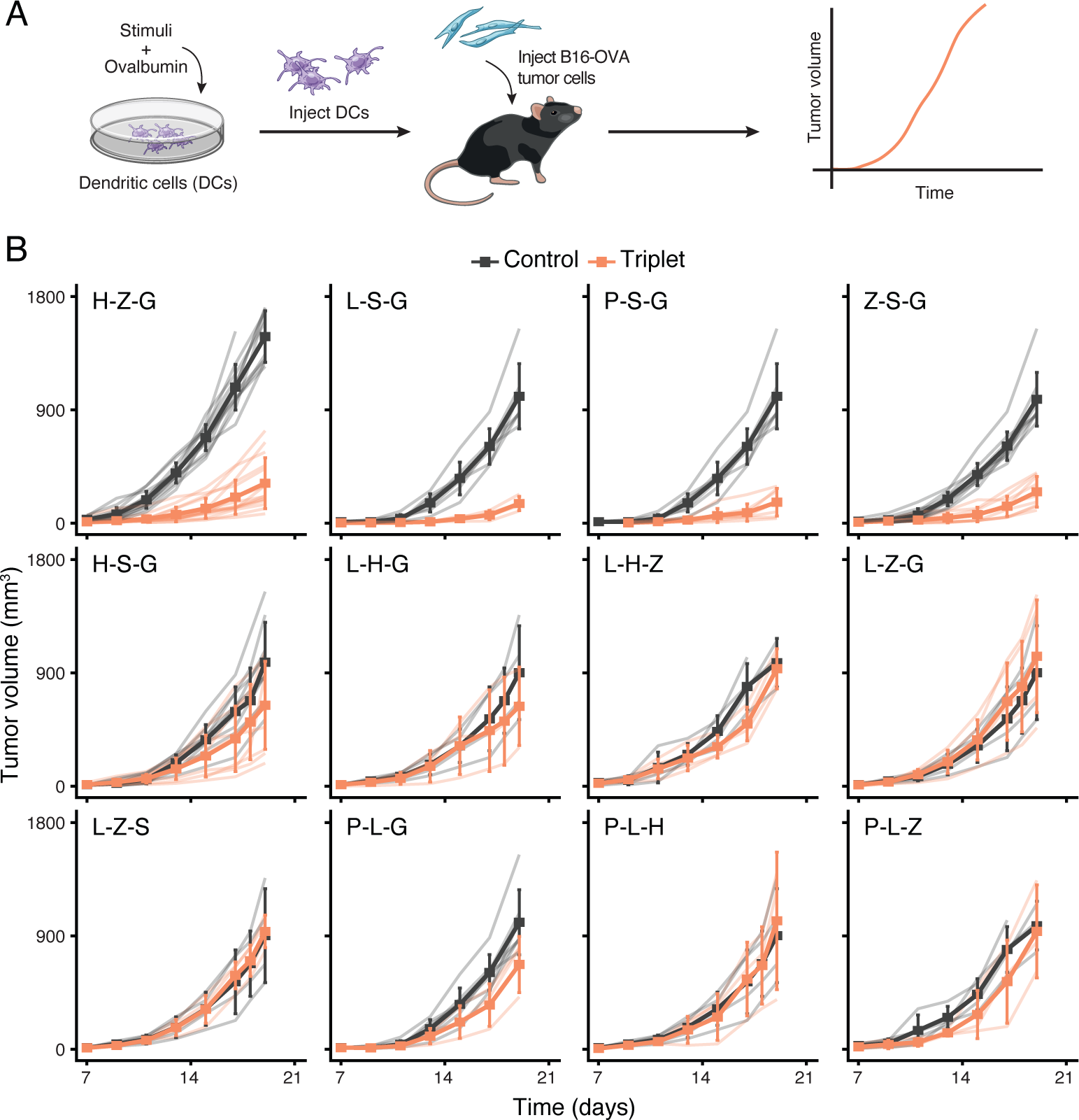
Adjuvant triplets generate potent cell-based vaccines against melanoma in mice. (A) Schematic overview of the experimental design. From left to right: mouse DCs are stimulated with adjuvant triplets and pulsed with ovalbumin (OVA) *in vitro*; then DCs are injected subcutaneously in mice which received 10^5^ OVA-expressing B16.F10 melanoma cells (B16-OVA) in the contralateral skin region; and tumor growth is measured over time to assess vaccine effects. (B) Average tumor growth (solid line) in cohorts of mice treated with DCs pulsed with OVA and stimulated with indicated adjuvant triplets (orange) or left unstimulated as control (black). Light color lines indicate the growth from each mouse within each cohort. Error bars, SEM (*n* = 3-15 mice per cohort).

Next, we sought to characterize the anti-tumor effects of the H-Z-G adjuvant triplet. First, we used tumor-bearing animals which were injected with OVA-loaded DCs that did not receive any stimulation and which showed similar tumor progression as mice inoculated with B16-OVA cells only (**Figure 6A**), and thus confirmed the adjuvant-dependent effects on decreasing tumor growth. In addition, the anti-tumor effects triggered by H-Z-G-stimulated DCs loaded with OVA (1) were antigen-specific (**Figure 6B**), (2) required the presence of T cells (**Figure 6C-D**), and (3) were applicable to OVA-expressing MC38 colorectal adenocarcinoma and E.G7 lymphoma cells (**Figure 6E**). H-Z-G-based DC therapy led to an increase in CD8^+^ T cell infiltration in the tumor, as opposed to a DC vaccine prepared with P-L-H that showed no effect on tumor growth (**Figure 6F**). Second, we found that the anti-tumor effects of the triplet H-Z-G were mediated in part (1) by endogenous, migratory DCs not exposed to adjuvant signals but likely activated by cytokines released by DC-based therapy, as shown by the use of *Batf3^-/-^* mice lacking conventional DC1 cells (Guilliams et al., 2014), and (2) by direct antigen presentation by the DC vaccine itself as shown by the decrease in anti-tumor activity when using DCs lacking MHC class II (**Figure 6G**). Third, the anti-tumor adjuvant triplet H-Z-G displayed stronger effects than those observed with its corresponding adjuvant singles and pairs (**Figure 6H**), and with a DC maturation cocktail used in a human DC vaccine against melanoma (Carreno et al., 2013; Carreno et al., 2015) (**Figure 6I**).

**Figure 6.**
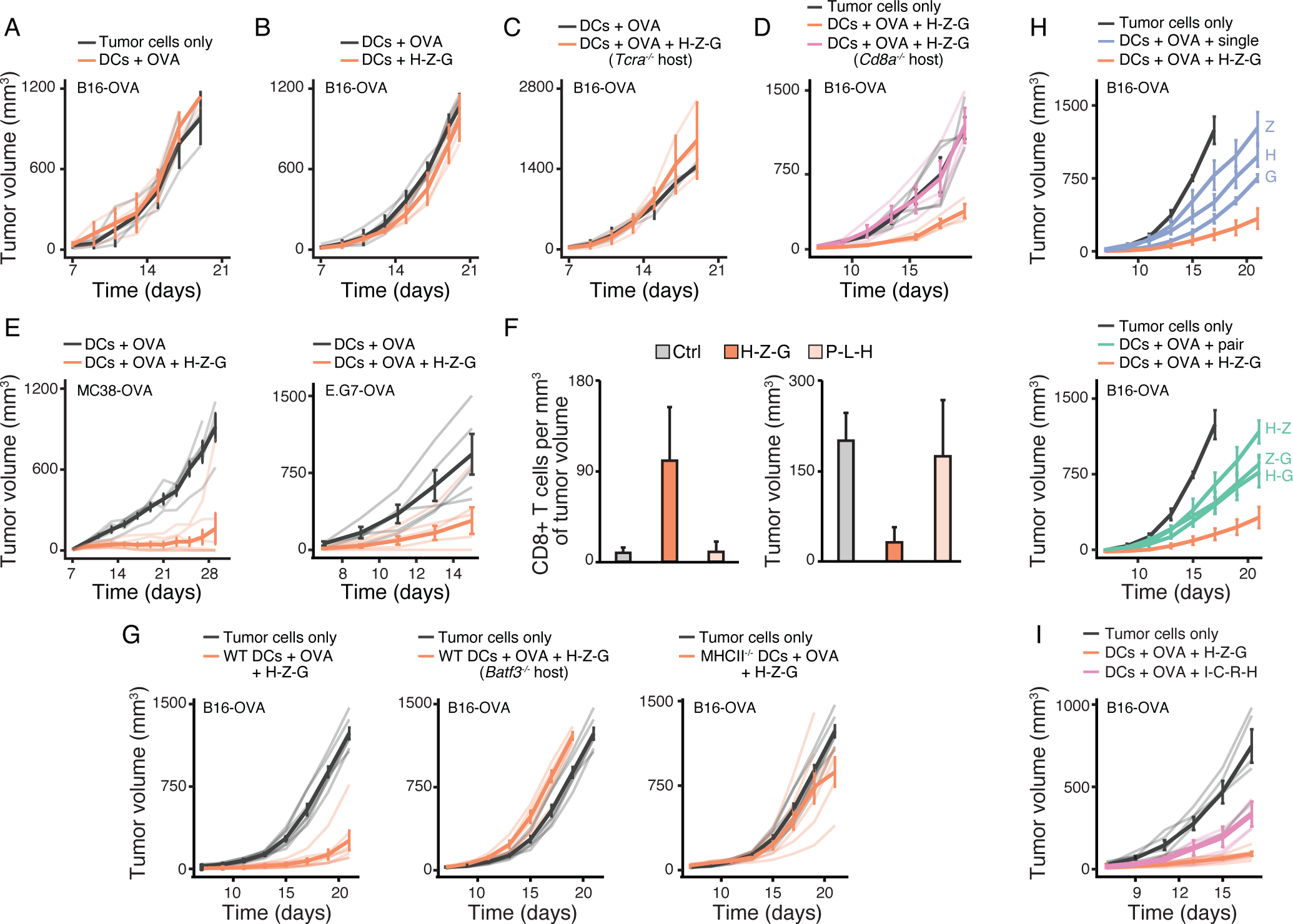
In vivo characterization of the anti-tumor effects mediated by the H-Z-G (poly(I:C)-Zymosan-cGAMP) adjuvant triplet using DC-based therapy. (A-E) Mean tumor growth (solid lines) in cohorts of wild-type mice, unless indicated otherwise for knockout strains (*Tcra^-/-^*, C; *Cd8a^-/-^*, D), injected with 10^5^ B16-OVA (A-D), 10^5^ MC38-OVA or 5 x 10^5^ E.G7-OVA (E) tumor cells and indicated DC vaccines. DCs + OVA, unstimulated DCs pulsed with ovalbumin protein (OVA); DCs + H-Z-G, DCs stimulated with the H-Z-G (poly(I:C)-Zymosan-cGAMP) ligand triplet without OVA; DCs + OVA + H-Z-G, DCs stimulated with H-Z-G and pulsed with OVA. Light color lines indicate tumor growth for individual mice within each cohort. Error bars, SEM (*n* = 3-6 mice per cohort). (F) Quantification of CD8^+^ T cells infiltrated in B16-OVA tumors (left) and corresponding tumor volumes (right) at day 13 post-injection of tumor cells and indicated DC vaccines. Ctrl, unstimulated DCs pulsed with OVA; H-Z-G and P-L-H, DCs pulsed with OVA and stimulated with ligand triplets H-Z-G or P-L-H (Pam3CSK4-LPS-poly(I:C)), respectively. Error bars, SEM (*n* = 3 mice per cohort). (G) Mean tumor growth (solid lines) in cohorts of wild-type mice (except for the group indicated as *Batf3^-/-^*, middle panel) injected with 10^5^ B16-OVA cells and left untreated as controls (tumor cells only; black lines), or treated (orange lines) as follows at day zero with wild-type (left and middle panels; WT) or MHC class II-deficient (right panel; MHCII^-/-^) DCs pulsed with OVA and stimulated with H-Z-G. Light color lines indicate tumor growth for individual mice within each cohort. Error bars, SEM (*n* = 4-7 mice per cohort). (H-I) Mean tumor growth (solid lines) in cohorts of wild-type mice injected with 10^5^ B16-OVA cells and left untreated as controls (tumor cells only; black lines), or treated with DCs pulsed with OVA and stimulated with indicated ligand singles (H, poly(I:C); Z, Zymosan; G, cGAMP), pairs (H-Z, H-G, Z-G) and triplet (H-Z-G) (H), or with the I-C-R-H quadruplet (IFN-γ at 100 U/mL, sCD40L at 100 ng/mL, R848 at 20 µg/mL, and poly(I:C) at 20 µg/mL) (I). Light color lines indicate tumor growth for individual mice within each cohort. Error bars, SEM (*n* = 4-9 mice per cohort).

### The effects of adjuvant triplets on tumors *in vivo* are explained by single and pairwise effects

Lastly, we asked whether the anti-tumor effects induced by the 4 out of 12 adjuvant triplets tested could be captured by the *in vivo* effects of single and pairwise stimulations of DC vaccines. To do so, we first sought to identify an *in vivo* proxy for the anti-tumor effects of the DC vaccination strategies discovered above. We measured the impact of the 12 DC vaccines tested in our B16-OVA model (**Figure 5B**) on the production of 12 T cell-associated cytokines in the inguinal draining lymph node (dLN) for the skin site of DC injection. A week after DC vaccine injection subcutaneously, the dLN was collected, dissociated and 5 x 10^5^ total dLN cells were kept in culture in the presence of the OVA protein (**Figure 7A, STAR Methods**). By PCA, the 12 triplets tested *in vivo* in our tumor model broadly clustered into three groups based on the cytokine profiles measured in the draining dLN, which revealed that the 4 triplets with anti-tumor effects separated from the other 8 (**Figure S6B, Table S6A**). We found that the levels of IL-17A production by dLN cells at 1 week post-vaccination were the most strongly anti-correlated with tumor volumes at ∼3 weeks post-tumor cell inoculation, suggesting that IL-17A could be used as a proxy for the anti-tumor effects of the DC vaccines used in this study (**Figure 7B** and **Figure S6A-C**). IL-6 and Il-17F also correlated well with anti-tumor effects but their levels were lower and closer to the detection limit of our assay (**Figure S6A**). Interestingly, IFN-γ levels were high in the 4 triplets leading to strong anti-tumor effects but were also found to be elevated in 3 out of the 8 triplets which did not lead to anti-tumor responses (**Figure S6A**).

**Figure 7.**
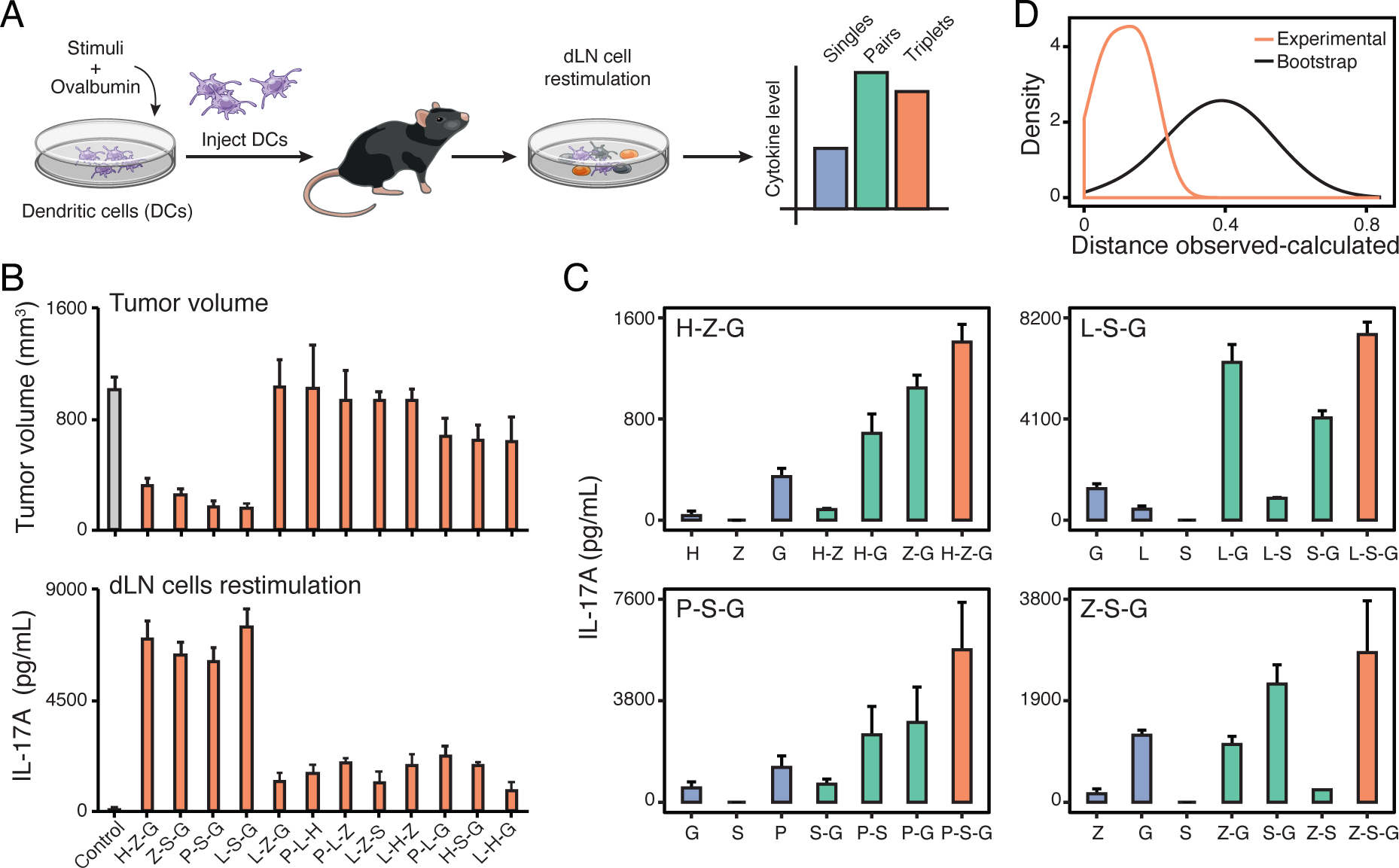
Adjuvant triplet effects on tumor growth are explained by singles and pairs. (A) Schematic overview of the experimental design. From left to right: mouse DCs are stimulated with adjuvant singles, pairs or triplets and pulsed with ovalbumin (OVA) *in vitro*. DCs are then injected subcutaneously in mice and a week later total draining lymph node (inguinal; dLN) cells are placed in culture with or without OVA. Cytokine concentrations are measured in the dLN cell culture supernatant after two days. (B) Relationship between tumor volume and IL-17A production in the dLN for indicated triplets used to prepare DC vaccines. Shown are B16-OVA tumor volumes at day 19 post tumor cell and DC vaccine injections (top), and IL-17A concentrations in dLN cell culture supernatants (bottom). Control, unstimulated DC vaccine pulsed with OVA only. Error bars, SEM (top, *n* = 4; bottom, *n* = 3-15 mice per cohort). (C) IL-17A production by inguinal dLN cells from mice injected subcutaneously with DC vaccines loaded with OVA and stimulated with indicated singles or combinations for the four adjuvant triplets with anti-tumor effects: H-Z-G (top left), L-S-G (top right), P-S-G (bottom left) and Z-S-G (bottom right). dLN cells were plated 7 days post-DC vaccination in medium containing 1 µg/mL of purified OVA protein. Blue, singles; green, pairs; orange, triplets. Error bars, SEM (*n* = 4). (D) Distribution of the distances between IL-17A concentrations for adjuvant triplets with anti-tumor effects (H-Z-G, L-S-G, Z-S-G and Z-S-G) that were experimentally observed and calculated using data from composite singles and pairs. Orange line, distribution from experimental values; black line, distribution observed by bootstrap analysis prior to calculating triplet values using Isserlis.

We found that IL-17A production by dLN cells was fully attributable to CD4+ T cells, as shown by cell depletion experiments (**Figure S6D**). In addition, using *Il17a^-/-^* or *Rorc^-/-^* mice abolished the anti-tumor effects of the adjuvant triplet H-Z-G (**Figure S6E-F**), which further reinforce the potential use of IL-17A production in the dLN as a surrogate for anti-tumor effects.

We then measured the levels of IL-17A production by dLN cells from mice injected with DC vaccines generated with the 4 triplets displaying anti-tumor effects, namely P-S-G, L-S-G, H-Z-G and Z-S-G, as well as their matching singles and pairs (**Figure 7C, Figure S6G-H, Table S6B**). Using the statistical framework established above for our in vitro DC-T system, we found that the levels of IL-17A induced by DC vaccines produced with adjuvant triplets were accurately described using only the data from DC vaccines made with the corresponding singles and pairs (**Figure 7D**). The correlation between observed and calculated IL-17A values was (1) maintained at two concentrations of OVA and without OVA being added to dLN cell cultures (**Figure 7C, Figure S6G-H**), and (2) statistically significant as shown by bootstrap analysis in which singles and pairs were scrambled prior to Isserlis calculations (**Figure 7D**). These results suggest that interactions among triplets of immune adjuvants are accurately described by single and pairwise effects at the level of the host.

## DISCUSSION

We studied the combinatorial effects of immune adjuvants using a representative set of microbial stimuli. Our data reveal an intrinsic property governing the combinatorial logic of microbial sensing: the effects of triplet combinations of adjuvants can be accurately described using only the effects of singles and pairs of stimuli. Remarkably, this simplifying property of pathogen-sensing pathways was applicable both in cell cocultures *in vitro* and in cell-based immunotherapies in mouse models of cancer. Our finding greatly simplifies the description of the combinatorial problem posed by the sensing of complex microbial or adjuvant inputs by innate immune pathways and their downstream impact on immunity. Overall, our findings are important for the fundamental understanding of how innate immunity processes information from complex inputs, which is critically important for the rational control of the innate immune system in therapy (Demaria et al., 2019).

What do our findings tell us about the properties of pathogen-sensing pathways in terms of their topology, information processing capacity and evolution? First, our results suggest that the wiring of the pathogen-sensing system allows pairwise interactions between pathways but limits, if not eliminate, higher-order interactions, which is reminiscent of results from disparate studies on other biological systems such as ecological and microbial interactions (Friedman et al., 2017; Grilli et al., 2017; Vandermeer, 1969), protein folding (Socolich et al., 2005), neuronal networks (Schneidman et al., 2006), and responses to antibiotics, drugs or agonists (Chatterjee et al., 2010; Wood et al., 2012; Zimmer et al., 2016). It remains unknown whether such inter-pathway wiring is similar in the context of interactions between pathogen-sensing and other pathways for stress, costimulatory or cytokine signals. Previous work by others combining one PRR agonist, LPS, and other non-PRR agonists found limited higher-order effects at the level of secretion of few cytokines (Hsueh et al., 2009), although it remains to be tested systematically using a variety of stimuli as inputs and by monitoring the net effects of stimuli combinations on immune responses as opposed to cytokine secretion. Interestingly, pioneering work by others on how T cells respond to combinatorial costimulatory and cytokine stimuli revealed an even simpler picture from the one proposed here for pathogen sensing, whereby pathways acted independently (Gett and Hodgkin, 2000; Marchingo et al., 2014). Thus, conducting comparative studies of the combinatorics of various immune signaling pathways will undoubtedly yield critical information to decipher and manipulate these complex systems. Future work is also needed to decipher the molecular mechanisms underlying the numerous cases of pairwise synergy and antagonism observed in this study by, for example, building upon recent systematic efforts to understand pairwise crosstalk between TLR pathways (Lin et al., 2017).

Second, the computation performed by innate immune cells sensing multiple microbial stimuli aims to rapidly identify a microbial threat and respond appropriately (Janeway, 1989). By decreasing the likelihood of higher-order interactions, the pathogen-sensing system perhaps increases its ability to reliably perform its functions – detecting microbes and transmitting information to adaptive immune cells (Iwasaki and Medzhitov, 2015) – in the face of perturbations, such as virulence factors (Finlay and McFadden, 2006) or inborn genetic errors (Bousfiha et al., 2018).

Third, why are pathogen-sensing pathways limited in the number of ways that they can interact? From a network biology standpoint, having some degree of shared and interacting nodes within a network of pathways is a likely requirement for the simplifying property we describe for PRRs to emerge. Although the exact level of node sharing and interactions is not known across PRR pathways, two decades of work on pathogen sensing has identified many shared components and examples of interactions between pathways (Chevrier et al., 2011; Crozat et al., 2009; Kawai and Akira, 2011; Loo and Gale, 2011; Osorio and Reis e Sousa, 2011; Thaiss et al., 2016). However, it remains unknown what a network of pathways should look like in terms of node sharing and interacting for the simplifying rule put forth by our results to apply. In other words, what are the lower and upper limits on node sharing and interacting that are needed within a network responding to multiple inputs? Thus, there is a need for theoretical (*e.g.*, simulations) and experimental (*e.g.*, pathway- or network-level engineering) work on this topic. From an evolutionary standpoint, one possibility is that while PRR pathways have been selected for by coevolutionary mechanisms between hosts and pathogens, the degree of crosstalk between PRR pathways is limited by the general evolutionary forces that guide the assembly of complex biological systems. The appearance of a novel pathogen-sensing pathway, or any other pathway, during evolution should not compromise the existing intracellular wiring of the cell. As a result, evolving a new pathway that could generate many-body interactions is perhaps an unlikely scenario. More generally, the constraints imposed by the sequential assembly of evolved systems is a possible explanation as to why pairwise correlations can sometime capture the complexity of biological systems.

Our data show that triplet stimulations regulates mRNAs, accessible chromatin loci or secreted proteins which are also found to be regulated at the level of pairs and singles. Although rare, few exceptions to this observation were found. For example, the levels of the *Il10* and *Il33* genes appeared significantly different in triplet stimulations compared to matching singles and pairs (**Figure S4D-H**). Further investigations are needed to uncover the transcriptional mechanisms underlying this finding. In addition, the effects of combinatorial microbial inputs on other functions of innate cells, such as phagocytosis, chemotaxis or metabolism, as well as on T cells in our coculture system remain to be studied.

We demonstrated that DC-based vaccines which are prepared with higher-order combinations of adjuvants can induce potent anti-tumor responses. Going further, we found that triplet adjuvant effects can largely be explained by pairwise and single adjuvant effects, using the production of IL-17A in the draining lymph node as a proxy for tumoricidal effects. Our findings provide a first test for the applicability of our model on the collective behavior of the pathogen sensing system *in vivo*. These observations have several critical implications for therapy and future research avenues. First, while DC-based vaccines have shown limited efficacy (Kantoff et al., 2010), partly owing to suboptimal DC maturation conditions (Garg et al., 2017; Sabado et al., 2017), we suggest that improved formulations based on the rationale combination of adjuvants could broaden the scope of therapeutic applications for such cell-based immunotherapies (**Figure 6I**) (Carreno et al., 2013; Carreno et al., 2015). Second, IL-17A could be useful as a biomarker for the generation of potent anti-tumor T cell responses and thus used for the screening of candidate immunotherapies that rely on vaccination or other modalities. While an increase in the number of IL-17-producing cells in the tumor environment correlates with improved survival for patients with esophageal squamous cell carcinoma (Lv et al., 2011), further work is needed to assess its precise role in our DC vaccine model. Perhaps IL-17 plays an indirect role on the promotion of cytotoxic T cell responses in the lymph node through the induction of other cytokines such as IL-12 and IL-6, as shown in other contexts (Benchetrit et al., 2002; Qian et al., 2017).

Overall, our work provides a conceptual framework to decipher the general rules governing the combinatorial logic of innate immune signaling and to help building predictive models for combining adjuvants to build vaccines and immunotherapies.

## STAR METHODS

### Mice

Female C57BL/6J (stock 000664), B6.Cg-*Ptprc^a^ Pepc^b^* Tg(TcrLCMV)1Aox/Ppmj (SMARTA-1; stock 030450), B6.129S(C)-Batf3^tm1Kmm^/J (*Batf3* knockout; stock 013755), B6(Cg)-*Sting1*^tm1.2Camb^/J (Sting-1 knockout; stock 025805), B6.129S6-*Clec7a*^tm1Gdb^/J (Dectin-1 knockout; stock 012337), B6.129P2(SJL)-Myd88^tm1.1Defr^/J (Myd88 knockout, stock 009088), C57BL/6J-Ticam1Lps2/J (Trif knockout, stock 005037), B6.129-Mavs^tm1Zjc^/J (MAVS knockout, stock 008634), B6.Cg-Ifih1^tm1.1Cln^/J (Mda5 knockout, stock 015812), B6.129S2-H2dlAb1-Ea/J (MHC class II knockout; stock 003584), *Il17a*^tm1.1(icre)Stck^/J (IL-17a knockout; stock 016879), B6.129P2-*Rorc*^tm1Litt^/J (RORγt knockout; stock 007571) and B6.129S2-*Cd8a*^tm1Mak^/J (CD8a knockout; 002665) mice were obtained from the Jackson Laboratories. OT-II^+^ TCRα^-/-^CD45.1^+/+^(OT-II) mice were kindly provided by Arlene Sharpe (Harvard Medical School, Boston, USA). Animals were housed in specific pathogen free and BSL2 conditions at The University of Chicago, and all experiments were performed in accordance with the US National Institutes of Health Guide for the Care and Use of Laboratory Animals and approved by The University of Chicago Institutional Animal Care and Use Committee.

### Cells

Bone marrow-derived dendritic cells (BMDCs) were generated from 6- to 8-week old female mice. Bone marrow cells were collected from femora and tibiae and plated at 2 x10^6^ cells in 10-cm non-tissue culture treated petri dishes (Corning 351029) in 10 mL of complete RPMI medium containing RPMI-1640 medium (ThermoFisher Scientific 11875119) supplemented with 10% volume/volume (v/v) heat-inactivated fetal bovine serum (Seradigm 1400-500), L-glutamine (2 mM, Corning 25005CI), penicillin and streptomycin (Lonza Biowhittaker 17602E), MEM non-essential amino acids (Corning 25025CI), HEPES (10 mM, Corning 25-060-CI), sodium pyruvate (1 mM, Corning 25000CI), β-mercaptoethanol (55 μM, Fisher Scientific 21-985-023). Recombinant murine GM-CSF (15 ng/mL; Peprotech 315-03-100μg) was added to the complete RPMI medium. Cells were fed at day 2, 5 and 7 with 3 mL of complete RPMI medium containing GM-CSF. At day 8, non-adherent cells were collected by pipetting, centrifuged and resuspended in fresh complete RPMI medium without GM-CSF. Cells were plated in 100 µL of medium in non-tissue culture treated 96-well flat bottom plates (ThermoFisher Scientific 260860) and incubated overnight prior to stimulations as indicated.

To enrich MHC class II (MHC-II)-positive DCs from total BMDC cultures, non-adherent cells were collected at day 8 and positively selected using anti-MHC class II microbeads (Miltenyi Biotec 130-052-401).

To purify the three DC populations present in total BMDC cultures based on the following markers: CD11c, CD11b and MHC-II, we used FACS. CD11c^+^ cells were stained with the following fluorescently conjugated antibodies (Biolegend): anti-CD11b (M1/70; 101205), anti-CD11c (N418; 117309), anti-IA/IE (M5/114.15.2; 107629) and DAPI (live/dead marker). Cells were sorted into CD11b^+^MHC-II^hi^, CD11b^+^MHC-II^med^, CD11b^+^MHC-II^lo^ subpopulations using a BD FACSAria II instrument.

To generate human blood monocyte-derived dendritic cells (moDCs), human whole peripheral blood from three healthy donors was obtained from Stemcell Technologies (catalog number 70504.5). Peripheral blood mononuclear cells (PBMCs) were isolated by density gradient centrifugation as follows: 15 mL of lymphocyte separation medium (LSM) (Corning 25072CV) was carefully underlaid beneath 35 mL of blood diluted 2X with 1X PBS/2 mM EDTA in a 50 mL tube and centrifuged at 400 *g* for 30 minutes at 20°C without break. The upper layer was aspirated leaving the white buffy coat interphase containing PBMCs undisturbed. The interphase was transferred to a new 50 mL tube containing 10 mL of 1X PBS, mixed and centrifuged at 400 *g* for 10 minutes at 20°C. Following the washing step, the supernatant was removed and PBMCs resuspended in 1X PBS. CD14^+^ Monocytes were then purified from total PBMCs using the Monocyte Isolation kit II (purity >90%; Miltenyi Biotec 130-091-153). CD14^+^ monocytes were plated at 10^6^ cells/mL in 10-cm non-tissue culture treated petri in 10 mL of complete RPMI medium prepared as above but without sodium pyruvate and MEM non-essential amino acids and by adding recombinant human IL-4 (40 ng/mL; Peprotech 200-04μg) and human GM-CSF (100 ng/mL; Peprotech 300-03μg). Cells were fed at day 3 and 5 with 5 ml of complete RPMI medium for human DCs supplemented with GM-CSF as described above. At day 7, non-adherent cells were collected by pipetting, centrifuged and resuspended in fresh complete RPMI medium without GM-CSF and IL-4. Cells were plated in 100 µL of medium in non-tissue culture treated 96-well flat bottom plates and incubated overnight prior to stimulations as indicated.

Ovalbumin-expressing B16.F10 (B16-OVA; a gift from Arlene Sharpe, Harvard Medical School, Boston, USA) and MC38 (MC38-OVA; a gift from Darrell Irvine, MIT, Cambridge, USA) cell lines were cultured in DMEM (ThermoFisher Scientific 11995073) supplemented with 10% v/v heat-inactivated FBS (Seradigm 1400-500) and penicillin/streptomycin (100 U/mL/100μg/mL, Lonza Biowhittaker 17602E). Ovalbumin-expressing E.G7 (E.G7-OVA; a gift from Darrell Irvine, MIT, Cambridge, USA) cells were cultured in complete RPMI medium.

Mouse splenic CD4^+^ T cells were isolated from OT-II or SMARTA-1 mice using CD4 (L3T4) MicroBeads (130-117-043) and LS columns (130-042-401) from Miltenyi Biotec and labeled with 1 µM CFSE (ThermoFisher Scientific C34554).

### Reagents

Lipopolysaccharide (L) from E. coli K12 (tlrl-peklps), Pam3CSK4 (P) (tlrl-pms), high-molecular weight polyinosinic-polycytidylic acid (poly(I:C) or H) (tlrl-pic-5), class B CpG oligonucleotide (ODN 1668 or C) (tlrl-1668-1), cyclic [G(3’,5’)pA(3’,5’)p] (3’3’-cGAMP or G for stimulating mouse cells) (tlrl-nacga-1), cyclic [G(2’,5’)pA(3’,5’)p] (2’3’-cGAMP or G for stimulating human cells) (tlrl-nacga23), Zymosan depleted which is a S. cerevisiae cell wall preparation (tlrl-zyd), and R848 (tlrl-r848) were purchased from Invivogen. Purified EndoFit grade ovalbumin (OVA) was obtained from Invivogen (vac-pova). Sendai Virus (SeV) was obtained from ATCC (VR-907). Recombinant murine IFN-γ (315-05) and sCD40L (315-15) were purchased from Peprotech. The antibody anti-CD4 was purchased from Biolegend (GK1.5; 100414), and DAPI from Biotium (40043).

### *In vitro* DC-T coculture

Mouse BMDCs prepared and plated in 96-well plates as detailed above (10,000 cells/well after overnight incubation) were incubated for 6 hours at 37°C with 200 µg/mL of OVA (for OT-II CD4^+^ T cells) or 0.005 μg/ml GP_61-80_ peptide (for SMARTA-1 CD4^+^ T cells) and with or without indicated ligands used alone or in combination at the following concentrations unless otherwise indicated: LPS, 100 ng/mL; PAM3CSK4, 250 ng/mL; 3’3’cGAMP or 2’3’cGAMP, 20 μg/mL; Zymosan depleted, 30 μg/mL; Sendai Virus, MOI 10; ODN 1668 CpG-B, 10 μg/mL; poly(I:C), 20μg/mL. After incubation, BMDCs were washed by medium replacement and fresh complete RPMI medium (100 µL) was added to the cultures. 50,000 freshly isolated and CFSE-labeled transgenic OT-II or SMARTA-1 T cells were then added to the DC culture in 100 µL of complete RPMI medium. Cocultures were incubated at 37°C for 3 days and T cells were harvested by pipetting and centrifugation. T cells were resuspended in 1X PBS buffer supplemented with 0.5% FBS and 2 mM EDTA (VWR BDH7830-1) and stained using an anti-CD4 antibody and DAPI to exclude dead cells. Flow cytometry data were acquired on the NovoCyte flow cytometer (Acea Biosciences/Agilent) and analyzed using the FlowJo software (BD).

For experiments whereby DCs stimulated with single ligands were mixed prior to coculture with T cells, mouse 100,000 DCs were incubated for 6 hours at 37°C with 200 µg/mL of OVA with single ligands. Supernatant was aspirated and 100 µL of PBS containing 10 mM EDTA was added to cells for 10 minutes at 37°C to detach them. Cells were harvested by pipetting, counted and plated at a 1:1 ratio (5,000 cells for each single ligand). 50,000 freshly isolated and CFSE-labeled transgenic OT-II T cells were then added to the DC culture in 100 µL of complete RPMI medium.

### Mouse tumor models

The abdomen of mice used for experiments were shaved using a pet trimmer (Wahl Bravmini CLP-41590) on the day before tumor cell and BMDC injections. Mice were injected subcutaneously with 500,000 E.G7-OVA, 100,000 B16-OVA or MC38-OVA cells resuspended in 100 µL of sterile saline in the flank. At the same time, mice were injected subcutaneously in the contralateral flank with 250,000 BMDCs that had been incubated with 25 µg/mL of OVA and indicated ligands for 6 hours at 37°C and washed three times with 1X PBS prior to being resuspended at 250,000 cells/100 µL in sterile saline. For consistency across experiments, tumor cells were thawed from liquid nitrogen stocks frozen in 90% FBS and 10% DMSO 2 days prior injections and passaged twice in total. Tumor volumes were calculated using the formula 1/2 × *D* × *d*^2^, where *D* is the major axis and *d* the minor axis (in mm). Mice were sacrificed when tumors reached 1000 cm^3^ or upon ulceration.

For analysis of tumor-infiltrating lymphocytes, B16-OVA tumors were dissected from mice, weighed and mechanically disaggregated before digestion with collagenase type I (400 U/ml; Worthington Biochemical) for 30 min at 37°C. After digestion, tumors were passed through 70-µm filters and lymphocytes were enriched by centrifugation using a gradient of 40/70% Percoll PLUS (GE Healthcare Life Sciences 17-5445-02). Cells were stained with the following fluorescently conjugated antibodies (Biolegend): anti-CD3e (145-2c11), anti-CD45 (30-F11), anti-CD8a (53-6.7) and Zombie-NIR (live/dead marker). Flow cytometry data were acquired on the NovoCyte flow cytometer (Acea Biosciences/Agilent) and analyzed using the FlowJo software (BD).

### Restimulation of total lymph node cells with ovalbumin

Mice were injected subcutaneously in both flanks with 250,000 BMDCs that had been incubated with 25 µg/mL of OVA and indicated ligands for 6 hours at 37°C and washed three times with 1X PBS prior to being resuspended at 250K cells/100 µL in sterile saline. Seven days after DC injections, the inguinal draining lymph nodes (dLNs) were collected from each mouse and minced using a microtube pestle (USA Scientific 1415-5390) in 1.5 mL tubes containing 500 µL of complete RPMI medium. Cell suspensions were filtered on a 100-µm filter mesh, centrifuged and resuspended in complete RPMI medium prior to counting cell concentrations. Cells were then plated at 500,000 total dLN cells/well in 200 µL of complete RPMI medium in non-tissue culture treated flat bottom 96-well plates (Thermofisher Scientific 260860). Cells were incubated with 1 or 10 µg/mL OVA or left untreated for 2 days at 37°C and 5% CO_2_. Cell culture supernatants were collected and stored in single-use aliquots at -80°C until further processing for measuring cytokine concentrations as described below.

For the culture of CD4+ T cells depleted dLN cells, CD4+ T cells were depleted from whole lymph node single cell suspension by positive selection using the MojoSort Mouse CD4 T Cell Isolation Kit (BioLegend 480006).

### Cytokine quantifications

Cell culture supernatants were collected from (1) BMDC (100,000 cells/well) cultures 8 hours after stimulation, (2) DC-T cocultures after a 3-day incubation period, or (3) total draining lymph node (dLN) cell cultures (500,000 cells/well) kept in culture for 2 days, and stored frozen at -80°C in single-use aliquots.

For sandwich Enzyme-Linked Immunosorbent Assay (ELISA), cell culture supernatants were diluted using the ELISA assay diluent (Biolegend 4212013) and cytokine concentrations were measured using the ELISA MAX standard set mouse IFN-γ (BioLegend 430801) and IL-17A (BioLegend 432501) kits according to the manufacturer’s instructions.

For flow cytometric, bead-based immunoassays, DC and dLN cell culture supernatants were diluted and processed using the LEGENDplex mouse anti-virus response panel (BioLegend 740622) and the LEGENDplex mouse Th cytokine panel (BioLegend 740740) kits, respectively. Data were acquired on the NovoCyte flow cytometer (Acea Biosciences/Agilent) and analyzed using the LEGENDplex software v8 (BioLegend).

### Secretome analysis

Mouse BMDCs (10^5^ cells/96-well in 100 µL of medium) were stimulated for 8 hours at 37°C in complete RPMI medium made with RPMI-1640 without phenol red (ThermoFisher Scientific 11835-030) and without FBS (serum-free conditions). Cell culture supernatants were collected by pooling 3 wells per conditions for a total volume of ∼330-390 µL in a 1.5-mL tube and centrifuged at 1000 *g* for 5 min to remove remaining cells. Supernatants were transferred to new tubes and centrifuged at 20,000 *g* for 10 min to remove cell debris.

Supernatants (∼300 µL) were denatured by adding 100 µL of 8 M urea (Sigma Aldrich U4883) and incubating for 5 min at room temperature (RT) with shaking at 800 rpm. Proteins were reduced with 5 mM dithiotreitol (Thermo Fisher scientific 20291) for 30 minutes at RT and alkylated with 10 mM iodoacetamide (Sigma A3221-1VL) for 30 minutes at RT in the dark with shaking at 1000 rpm. Proteins were digested with 0.5 µg of trypsin (Promega V5113) for 16 h at room temperature with shaking at 700 rpm. The digestion was stopped by acidification by adding 4µL of formic acid (Honeywell Fluka 56302-10X1ML) to obtain a pH < 3 (pH indicator strips, EMD 9586). Peptide samples were desalted on C18 stage tips and to enable multiplexing, peptide samples were labeled with TMT-10 reagents (Thermo Scientific). The TMT-labeled samples were loaded on C18 stage tips and separated into 6 high-pH fractions using elution solvents containing ammonium formate buffer (0.0175% NH4OH, Sigma-Aldrich; 0.01125% formic acid, Fluka; 2% acetonitrile, Honeywell) and 10, 15, 20, 22.5, 25 and 50 % acetonitrile (Honeywell).

Tryptic peptides were analyzed on an EASY-nLC 1200 system coupled to a Q-Exactive Plus (ThermoFisher Scientific). The EASY-nLC system was equipped with a 75 µm x 20 cm column (packed in-house with 1.9 um C18 resin; Reprosil Gold, Dr. Maisch) and operated at a flow rate of 250 nL/min applying a 110 min linear gradient from 2 to 90 % solvent B (90 % ACN, 0.1 % FA) in A (3 % ACN, 0.1 % FA). MS measurements were performed on Q Exactive Plus with the following modifications: MS1 spectra were recorded at a resolution of 60k using a maxIT of 10 ms. Fragment spectra were acquired at 45k resolution using a maxIT of 86 ms for proteome measurements.

### RNA extraction

Mouse BMDCs (100,000/96-well or 10,000/96-well for sorted BMDCs) and human moDCs (10,000/96-well) were stimulated with indicated ligands for 6 hours at 37°C, and after stimulation culture supernatants were removed by aspiration. Cells were lysed with 30 µL of RLT buffer (Qiagen 79216) containing 1% (v/v) β-mercaptoethanol (VWR 97604-848). Total RNA was isolated using Dynabeads MyOne Silane (Thermo Fisher Scientific 37002D) and RLT buffer using a custom protocol(Kadoki et al., 2017). Remaining genomic DNA was removed by on-bead DNase I (Thermo Fisher Scientific AM2239) treatment at 37°C for 20 min. After washing two times with 80% ethanol, total RNA was eluted from beads in nuclease-free water.

### RT-qPCR

Total RNA was reverse transcribed using the High Capacity cDNA Reverse Transcription Kit (ThermoFisher Scientific 4368813) with both random nonamers (N9) and oligo(dT) primers. Real-time quantitative PCR reactions were performed on the CFX384 Real-Time PCR Detection System (Bio-Rad Laboratories) with LightCycler 480 SYBR Green I Master mix (Roche) and 0.5 mM of each primer in a final volume of 10 µL with 40 cycles of denaturation at 95°C for 15 s and annealing/extension at 60°C for 40 s. The following forward-reverse primer pairs were used to measure levels the following mouse genes: *Gapdh* (5’-ggcaaattcaacggcacagt-3’, 5’-agatggtgatgggcttccc-3’), *Tlr2* (5’-aagaggaagcccaagaaagc-3’, 5’-cgatggaatcgatgatgttg-3’), *Tlr3* (5’-cacaggctgagcagtttgaa-3’, 5’-tttcggcttcttttgatgct-3’), *Tlr4* (5’-acctggctggtttacacgtc-3’, 5’-ctgccagagacattgcagaa-3’), *Tlr9* (5’-actgagcacccctgcttcta-3’, 5’-agattagtcagcggcaggaa-3’), *Ddx58* (5’-ccacctacatcctcagctacatga-3’, 5’-tgggcccttgttgttcttct-3’), *Tmem173* (5’-tgaaaggctcttcattgtctctt-3’, 5’-tggcatcttctgcttcctaga-3’), *Clec7a* (5’-atcagcattcttccccaactcg-3’,5’-cagttccttctcacagatactgtatga-3’), *Cxcl10* (5’-gccgtcattttctgcctca-3’, 5’-cgtccttgcgagagggatc-3’), *Tnf* (5’-ccctcacactcagatcatcttct-3’, 5’-gctacgacgtgggctacag-3’), *Cxcl1* (5’-ctgggattcacctcaagaacatc-3’, 5’-cagggtcaaggcaagcctc-3’), *Ifnβ* (5’-ctggcttccatcatgaacaa-3’, 5’-agagggctgtggtggagaa-3’), *Il6* (5’-tgttctctgggaaatcgtgga-3’, 5’-gctacgacgtgggctacag-3’) and human gene: *GAPDH* (5’-agccacatcgctcagacac-3’, 5’-aatacgaccaaatccgttgact-3’). Amplification products were subjected to melting curve analysis using the CFX Manager System (Bio-Rad Laboratories) to exclude the amplification of non-specific products.

### RNA-seq

Multiplexed RNA-seq libraries were prepared using the following overall workflow(Kadoki et al., 2017): (1) oligo(dT)-primed RT reaction with sample barcoding followed by cDNA pooling; (2) single-primer PCR amplification; and (3) full-length cDNA tagmentation and amplification by PCR.

First, total RNA samples obtained from 1x10^5^ mouse BMDCs or 1x10^4^ human moDCs were reverse transcribed to cDNA by denaturing 10-µL RNA samples with 1 μL containing 2 pmoles of a custom RT primer, which is biotinylated in 5’ and containing sequences from 5’ to 3’ for the Illumina read 1 primer, a 6-bp cell barcode, a 10-bp unique molecular identifier (UMI) and an anchored oligo(dT)30 for priming (5’-/5Biosg/ACACTCTTTCCCTACACGACGCTCTTCCGATCT[6-bp barcode]NNNNNNNNNNTTTTTTTTTTTTTTTTTTTTTTTTTTTTTTVN-3’; where 5Biosg =5’biotinylation, V=A, G or C, N=A, G, C or T), at 72°C for 2 min and snap cooled on ice. A 9-µL RT mix containing 4 μL of 5X RT buffer, 1 μL of 10 mM dNTPs, 2 pmoles of template switching oligo (5’-iCiGiCACACTCTTTCCCTACACGACGCrGrGrG-3’; where iC = iso-dC, iG = iso-dC, rG = RNA G), 0.5 µL Maxima H Minus Reverse Transcriptase (ThermoFisher Scientific EP0753) and 3.5 µL of nuclease-free water was added to the denatured RNA samples, and plates were incubated at 42°C for 120 min. Next, double-stranded cDNA samples were pooled using DNA Clean & Concentrator-5 columns (Zymo Research D4013), and residual RT primers were removed using exonuclease I (New England Biolabs M0293).

Second, pooled full-length cDNA was amplified with 4-6 cycles of single-primer PCR using the following primer: 5’-/5Biosg/ACACTCTTTCCCTACACGACGC-3’ (5Biosg = 5’ biotinylation) and the Advantage 2 PCR Kit (Clontech 639206) in a 50-µL reaction volume and using the following cycling condition: 1 cycle at 95°C for 1 min; 4-6 cycles at 95°C for 15 sec, 65°C for 30 sec, 68°C for 6 min; and 1 cycle at 72°C for 10 min. Amplified cDNAs were cleaned up using 0.6X volume of magnetic beads Agencourt AMPure XP (Beckman Coulter A63880) and quantified using the Qubit dsDNA High Sensitivity Assay Kit (ThermoFisher Scientific Q32851). Third, 1 ng of cDNA was tagmented and amplified by PCR using the following forward forward 5’-aatgatacggcgaccaccgagatctacactctttccctacacgacgctcttccg*a*t*c*t-3’, where * indicates phosphorothioated DNA bases, and Illumina i7 reverse primers using the Nextera XT Kit (Illumina) with the following cycling conditions: 1 cycle at 72°C for 3 min; 1 cycle at 95°C for 30 sec; 12 cycles at 95°C for 10 sec, 55°C for 30 sec and 72°C for 30 sec; and 1 cycle at 72°C for 5 min. Libraries were cleaned up using 0.8X volume of magnetic beads Agencourt AMPure XP and gel purified using E-Gel EX Agarose Gels, 2% (ThermoFisher Scientific G402002), quantified with the Qubit dsDNA High Sensitivity Assay Kit (ThermoFisher Scientific Q32851), and sequenced on the NextSeq550 platform (Illumina) using the NextSeq 500/550 high output kit v2 and following sequencing conditions: 17 cycles for Read 1, 8 cycles for Index 1, 66 cycles for Read 2.

### Chromatin accessibility

The original ATAC-seq protocol(Buenrostro et al., 2013) was used with modifications as follows. 50,000 mouse BMDCs were centrifuged and cell pellets were lysed in 50 µL of ice-cold lysis buffer containing 10 mM Tris-HCl, pH 7.4, 10 mM NaCl, 3 mM MgCl_2_ and 0.1% IGEPAL CA-630, and immediately centrifuged at 500 g for 10 min at 4°C. Supernatants were discarded and pelleted nuclei were processed for tagmentation by adding the following mix: 22.5 µL of nuclease-free water, 2.5 µL of transposase and 25 µL of TD buffer from the Nextera DNA library prep kit (Illumina FC-121-1030), and by incubating the mixture at 37°C for 30 min. Tagmented genomic DNA was purified using DNA Clean & Concentrator-25 columns (Zymo research D4033). Sequencing libraries were generated using the following forward (5’-aatgatacggcgaccaccgagatctacactcgtcggcagcgtcagatgtg-3’) and barcoded reverse (5’-caagcagaagacggcatacgagat[8-bp barcode]gtctcgtgggctcggagatgt-3’) primers, and by performing 12 cycles of amplification with the Q5 Hot Start High-Fidelity 2X Master Mix (New England Biolabs M0494) using the following cycling conditions: 1 cycle at 72°C for 5 min; 1 cycle at 98°C for 30 sec; and 12 cycles at 98°C for 10 sec, 63°C for 30 sec and 72°C for 1 min. Libraries were purified using DNA Clean & Concentrator-25 columns to remove remaining primers, and amplicon size distributions measured using high sensitivity D5000 screentape (Agilent Technologies 5067-5592). Libraries were then quantified using the Qubit dsDNA High Sensitivity Assay Kit (ThermoFisher Scientific Q32851) and sequenced on the NextSeq550 platform (Illumina) using the NextSeq 500/550 high output kit v2 and following sequencing conditions: 42 cycles for Read 1, 8 cycles for Index 1, 42 cycles for Read 2.

### Analysis of flow cytometry data on T cell proliferation

The ultimate goal of our analysis is to test whether the effects of single and pairwise microbial inputs (*i.e.*, ligands for pattern-recognition receptors) can help to explain the effects of higher-order combinations (*i.e.*, triplets of ligands).

We describe below all of the main steps of our analysis and the full code is publicly available in the following repository: https://github.com/chevrierlab/combos-paper.

(1) *CFSE data processing:* We analyzed raw flow cytometric data using the FlowJo software to calculate live and dead CD4^+^ T cells. Representative plots for our gating strategy are shown in Figure S1. Next, we manually drew gates for each CFSE peak in a given profile to calculate the numbers of T cells per division (referred to as cell generation).
(2) *Computing statistics of CFSE profiles:* To characterize the cellular proliferation profiles, we computed commonly used statistics for CFSE measurements (Roederer, 2011). The percent of T cells in the final population that have divided (proportion of divided T cells) was calculated by dividing the number of cells present in all the peaks below the undivided peak (*i.e.*, cell generation 0) by the total number of live cells. Next, we calculated the following metrics: the number of cells present at the start of the coculture (starting cells), given by 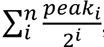,

i. the number of activated cells (cells that went into division) is the number of cells present at the start (*i*) minus the number of cells which did not divide (*i.e.*, cells in peak zero),
ii. the total number of divisions, given by 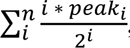,
iii. the total number of cells, given by 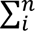, and
iv. the total number of cells which divided at least once is the total number of cells
v. minus the number of cells which did not divide (i.e., cells in peak zero).

Using the metrics listed above, we calculated the following six metrics that are commonly used to characterize CFSE proliferation profiles^32^:

a. The proliferation index (*Pi*) is the total number of divisions (*iii*) divided by the number of activated cells (*ii*), which provides the average number of divisions undergone per activated cell.
b. The division index (*Di*) is the total number of divisions (*iii*) divided by the number of starting cells (*i*), which provides the average number of divisions per starting cell.
c. The precursor frequency (*Pf*) is the number of activated cells (*ii*) divided by the number of starting cells (*i*), which is the probability that a cell will divide at least once.
d. The expansion index (*Ei*) is the total number of cells in the culture (*iv*) divided by the number of cells at the start (*i*), which is the fold expansion over the culture time.
e. The replication index (*Ri*) is the total number of divided cells (*v*) divided by the number of activated cells (ii), which is the fold expansion for activated cells.
f. The fraction diluted (*Dil*) is the total number of divided cells (*v*) divided by the total number of cells (*iv*), which is the fraction of cells in the final culture which divided at least once.

In addition, we normalized experimental *Pi*, *Di*, *Pf*, *Ei*, *Ri* and *Dil* values across all singles (7), pairs (21) and triplets (35) by dividing each value by the maximum value measured within each technical replicate of each experiment. Resulting normalized indices were used for computations of interaction scores and Isserlis calculations as delineated below.

(3) *Computing interaction scores for pairs and triplets:* To characterize the interactions that emerge when two ligands are combined (*i.e.*, two PRR pathways being activated), we compute a pairwise interaction score *I_AB_* for two ligands *A* and *B* and given by
 

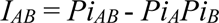

where *Pi* values are the proliferation indices computed as described above for both singles *a* and *b* and the pair *ab*. If *I_ab_* is close to zero, then *Pi_AB_* = *Pi_A_Pi_B_*, which is equivalent to Bliss independence, a common phenomenological model used in pharmacology to describe non-interacting drug pairs(Bliss, 1939; Bliss, 1956). To define regions of approximately additive behavior between two ligands, we computed the mean of the standard errors of all 21 *Pi_AB_* values obtained, which reflects the compounded measurement error for our experimental assay. We used the mean of the standard errors *m*, shown as a dotted line in Figure 2B and Figure S4D, as a threshold to define ligand pairs with synergistic (*I_AB_* > +*m*), antagonistic (*I_AB_* < -*m*), and additive (-*m* < *I_AB_* < +*m*) behaviors.

The starting hypothesis of this analysis implies that the net effect of a ligand triplet combination arises from the cumulative effect of the pairwise interactions(Wood et al., 2012). To assess this, we calculated a triplet interaction score *I_ABC_* for three ligands *A*, *B* and *C* and given by

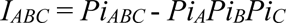

where *Pi* values are the proliferation indices computed as described above for the singles *A*, *B* and *C* and for the triplet *ABC*. *I_ABC_* provides a metric to measure the level of net pairwise and triplet interactions by subtracting single ligand effects from the triplet proliferation index. First, similar to above for pair interaction scores, we used the *I_ABC_* score to classify the interaction between three pairs as synergistic, antagonistic or additive, using the same thresholding approach based on the mean of the standard errors for *I_ABC_* values. Second, we compared *I_ABC_* values obtained through experimental measurements to those obtained by a statistical approach that uses only information from singles and pairs and defined below by an Isserlis formula.

(4) *Statistical modeling of triplet effects from single and pair effects:* Following upon the work of Wood and colleagues (Wood et al., 2012), we hypothesized from the outset that an equation derived from the Isserlis theorem (Isserlis, 1918), originally used to describe moment relationships, could serve as a statistical model to computer triplet effects using data from singles and pairs only. To test this, we used the following Isserlis formula:

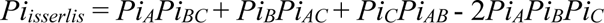

where *Pi* values are computed and normalized as explained above from experimental values. *Pi_isserlis_* values were computed from normalized *Pi* values for each of the 35 triplets and averaged across technical and biological replicate experiments. The correlation between average *Pi_isserlis_* values and experimentally observed triplet *Pi* values (*Pi_ABC_*) was measured by calculating an R-squared statistic using the lm() function in R (https://www.r-project.org/).

(5) *Bootstrap analysis:* We used bootstrapping to estimate the statistical significance of the correlation observed between averaged experimental (*Pi_ABC_*) and calculated (*Pi_isserlis_*) proliferation indices for triplets. We randomized normalized, experimental *Pi* values for singles and pairs across all experimental replicates generated in this study. *Pi_isserlis_* values were then computed from these randomized *Pi* data sets and compared to experimental *Pi_ABC_* values by calculating an R-squared statistic as above. We plotted the distribution of bootstrapped R-squared values from 500,000-1,000,000 randomizations. Lastly, we computed a *p*-value by calculating the fraction of bootstrapped R-squared values higher than the observed R-squared value.

### RNA sequencing data analysis

Sequencing read files were processed to generate raw (1) read and (2) UMI count matrices. For (1), we used the RNA-seq pipeline in the bcbio-nextgen project version 1.1.5 (https://bcbio-nextgen.readthedocs.io/en/latest/). Reads were aligned to the human hg19 genome or the mouse mm10 genome augmented with transcripts from Ensembl release 78 with STAR version 2.6.1d(Dobin et al., 2013). Quality control metrics were compiled with a combination of FastQC (http://www.bioinformatics.babraham.ac.uk/projects/fastqc/), Qualimap(Garcia-Alcalde et al., 2012), MultiQC (https://github.com/ewels/MultiQC) and custom metrics (3 million mapped reads were obtained on average per sample). Expression quantification was performed using both featureCounts version 1.4.4(Liao et al., 2014) with multi-mapping reads excluded and Sailfish version 0.9.2(Patro et al., 2014) with a kmer size of 31 with 30 bootstrap samples. For (2), we used custom scripts to map Read 2 sequences onto RefSeq mRNAs using BWA version 0.7.15(Li and Durbin, 2009), demultiplex the output based on barcodes stored in Read 1 (first 6 bp), and computed gene expression using UMIs stored in Read 1 (base 7 to 16) to produce raw UMI count matrices.

Differential expression (DE) analysis was done using custom scripts in R (https://www.r-project.org/). Raw count matrices were normalized across samples using the calcNormFactor function in edgeR(Robinson et al., 2010) and subsequently filtered to keep genes with at least 50 counts per million (cpm) in 2 samples. We identified DE genes using the following cutoffs: a 1.5-fold change with a Benjamini and Hochberg FDR adjusted *p*-value < 0.01 by comparing cells stimulated with each ligand combination to untreated, control cells using limma (http://bioinf.wehi.edu.au/limma/) (Ritchie et al., 2015).

To ask if a gene *x* found to be differentially regulated upon stimulation with a given triplet *ABC* is also regulated in any of the composite conditions of that triplet (i.e., singles *A*, *B*, *C* and pairs *AB*, *AC*, *BC*), we used the following criteria: (1) gene *x* is regulated in condition *ABC* compared to control cells using the same thresholds as above (fold-change > 1.5 with FDR < 0.01); (2) gene *x* is not regulated in the composite conditions of triplet *ABC* (*i.e.*, *A*, *B*, *C*, *AB*, *AC*, or *BC*) compared to control using the same thresholds as in (1); and (3) the level of gene *x* is significantly different between condition *abc* and all of its composite treatments (*i.e.*, *A*, *B*, *C*, *AB*, *AC* or *BC*) using as a threshold a Benjamini and Hochberg FDR adjusted *p*-value < 0.1. The number of genes which followed these three criteria were counted for each triplet, and the proportion of newly regulated genes for each of the 35 triplets was calculated as the ratio between the number of newly and all regulated genes.

To ask if a gene displayed a change in expression that is lower or higher upon triplet stimulation than that of the change observed in all matching stimuli singles and pairs, we computed an expression divergence metric for each gene regulated by a triplet by subtracting the log-fold change values of composite singles or pairs from the log-fold change of the corresponding triplet. The smallest value obtained between a given triplet and its matching composite singles and pairs was used as expression divergence.

### ATAC-seq data analysis

Sequencing reads were aligned to the mouse mm10 genome using bowtie2 version 2.2.9 (http://bowtie-bio.sourceforge.net/bowtie2/index.shtml) with the a maximum fragment length of 2000, and were sorted using samtools version 1.4.1 (http://samtools.sourceforge.net/). Peaks were called using MACS2 version 1.4 with a q-value threshold of 0.01 and a fixed background lambda using the following command: parameters callpeak --gsize 1.87e9 --nomodel -t out/${name}/${name}_sorted.bam -n out/${name}/${name} --nolambda --slocal 10000 -q .01

Peaks found in each sample were merged into a joint set of all peaks using the merge function in bedtools version 2.26(Quinlan and Hall, 2010). Reads for each peak were counted across experimental conditions using featureCounts() from the Rsubread package(Liao et al., 2019).

To identify differentially accessible peaks across conditions and newly regulated peaks in triplets compared to matching composite treatments, we used the same procedure as the one described above for RNA-seq. Peaks were considered differentially accessible in a treatment if they were different from the control by a fold-change greater than 1.5 and an FDR < 0.01.

### Principal component analysis of gene expression and chromatin accessibility data

Log2 fold change values between treated groups and the control group were obtained for each gene using limma, scaled to unit variance and centered by subtracting the mean before applying the prcomp() function in R. For human RNA-seq data analysis, PCA was performed on all three donors together and results were displayed for each donor individually.

### Secretome data analysis

For preprocessing of the data from peptide and TMT 10 experiments, peptide identification and quantification were performed using MaxQuant (version 1.6.0.1)(Cox and Mann, 2008). For the quantification at the MS/MS level, ’Reporter ion MS2’ was enabled and ’10plexTMT’ isobaric labels were selected. Tandem mass spectra were searched against the mouse reference proteome (mouse uniprot fasta) supplemented with common contaminants. For all searches carbamidomethylated cysteine was set as fixed modification and oxidation of methionine and N-terminal protein acetylation as variable modifications. Trypsin/P was specified as the proteolytic enzyme with up to 2 missed cleavage sites allowed. Results were adjusted to a 1% false discovery rate (FDR). The reporter-ion intensities were corrected for isotopic impurities before using the reporter-ion signals in each MS/MS spectrum for quantitative calculations.

For differential expression analysis, the corrected reporter ion intensities obtained from the mass-spectrometric measurements were divided by the internal standard mix reporter ion intensities and log2-transformed using custom scripts in R. The internal standard was hereby an equal mix of all analyzed samples. The log2 fold change for the treated vs. control samples were calculated, median-MAD normalized and analyzed for significant differences by a one-sample moderated T test. All identifications were considered significant with a Benjamini-Hochberg adj.p< 0.1.

For batch correction between experiment 1 (P-S-G triplet and composite ligand singles and pairs) and 2 (Z-S-G and P-L-H triplets and composite ligand singles and pairs), log2 fold change values calculated above were corrected using the ComBat() function from the sva package in R (http://bioconductor.org/packages/release/bioc/html/sva.html). Batch corrected log2 fold change values were averaged across replicates, then scaled to unit variance and centered by subtracting the mean before applying the prcomp() function in R for principal component analysis.

To benchmark the data obtained by our secretome analysis, we compared it to cytokine concentration values obtained by a bead-based immunoassay performed on supernatants from BMDC cultures and focusing on the following cytokines: CXCL1, TNF-α, CCL2, CCL5, CXCL10 and IL-6, which were detected by mass spectrometry and could be measured using a commercially available kit (LEGENDplex mouse anti-virus response panel; BioLegend 740622). Average log2 fold changes were computed between each treated group and the untreated control group. A linear model for the relationship between the log2 fold change values from secretome and immunoassay measurements was calculated using the lm() function in R.

### Generation of heatmaps

Heatmaps for RNA-seq, ATAC-seq and secretome data display the indicated numbers of transcripts, loci and proteins, respectively. Color intensities are determined by log2 fold change values for each heatmap. The rows of each heatmap were ordered by hierarchical clustering of log2 fold change values using one minus Pearson’s correlation as a distance metric. All heatmaps were generated using the Morpheus software (https://software.broadinstitute.org/morpheus/).

### Analysis of lymph node cell restimulation data

For principal component analysis, cytokine concentration values obtained by bead-based immunoassays were averaged within treatment and control groups, then scaled to unit variance and centered by subtracting the mean before applying the prcomp() function in R.

For statistical modeling of triplet effects from single and pair effects, IL-17A concentration values were averaged within each experiment, treatment group and OVA concentration. Average values were normalized by dividing each value by twice the maximum value within a given experiment. Normalized IL-17A concentration values were averaged across experiments for each treatment and OVA concentration group. We hypothesized that the same Isserlis statistical approach used above to calculate the proliferation index of T cells *in vitro* could be used to calculate IL-17A values, which serve as proxy for the effect of the T cell response on tumor growth. To do so, we subtracted normalized, averaged IL-17A concentration values, refered to as *Il*, from one and used the following formula:

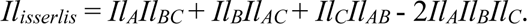

Next, to compare IL-17A values obtained experimentally for a given triplet (*Il_ABC_*) to those obtained with the Isserlis formula (*Il_isserlis_*), we plotted the distribution of the distances between these two values (*Il_isserlis_ – Il_ABC_*).

To estimate the significance of our Isserlis calculations, we used bootstrapping similarly to what is described above for the analysis of our *in vitro* coculture data. Here we randomized experimental IL-17A values (after averaging, normalization and subtracting from one) for singles and pairs across experiments prior to calculating *Ilisserlis* as above. The distribution of the *Il_isserlis_ – Il_ABC_* values was then plotted using bootstrap values from 1000 randomizations.

### Data availability

Sequencing and mass spectrometry data have been respectively deposited in the Gene Expression Omnibus under accession numbers GSE134869, GSE134874 and GSE134867, and in the MassIVE repository (http://massive.ucsd.edu). All scripts and preprocessed datasets are publicly available at the following repository: https://github.com/chevrierlab/combos-paper.

## ACKNOWLEDGEMENTS

We thank Peter Sage, Vikram Juneja, Hugo Carillon, and Motohiko Kadoki for help with pilot experiments; Andrew Ferguson, Melody Swartz, Savas Tay, Albert Bendelac, Jeffrey Hubbell, Peter Savage, Thomas Gajewski, Rama Ranganathan, Stefano Allesina, and Andrew Murray for valuable discussions; and Sigrid Knemeyer for help with artwork. This project was supported by an Institutional Research Grant (#IRG-16-222-56) from the American Cancer Society, the University of Chicago Medicine Comprehensive Cancer Center Support Grant (#P30 CA14599), an NIH Director’s New Innovator Award (1DP2AI145100-01), the Elliot and Ruth Sigal MRA Young Investigator Award and funds from the Bauer Fellows Program and the Pritzker School of Molecular Engineering (all to N.C.), and an NSF grant Physics of Living Systems (#129334) (to P.C.).

## AUTHOR CONTRIBUTIONS

Conceptualization, P.C. and N.C.; Methodology, S.P. and N.C.; Investigation, S.P. and N.C.; Mass spectrometry analysis, T.K. and P.M.; Isserlis analytical framework, P.C.; Formal analysis, A.G., P.C. and N.C.; Writing – Original Draft, S.P. and N.C.; Writing – Review & Editing, S.P., A.G., T.K., P.M., P.C. and N.C.; Funding Acquisition, P.C. and N.C.; Supervision, N.C.

## SUPPLEMENTARY FIGURE LEGENDS

**Figure S1.**
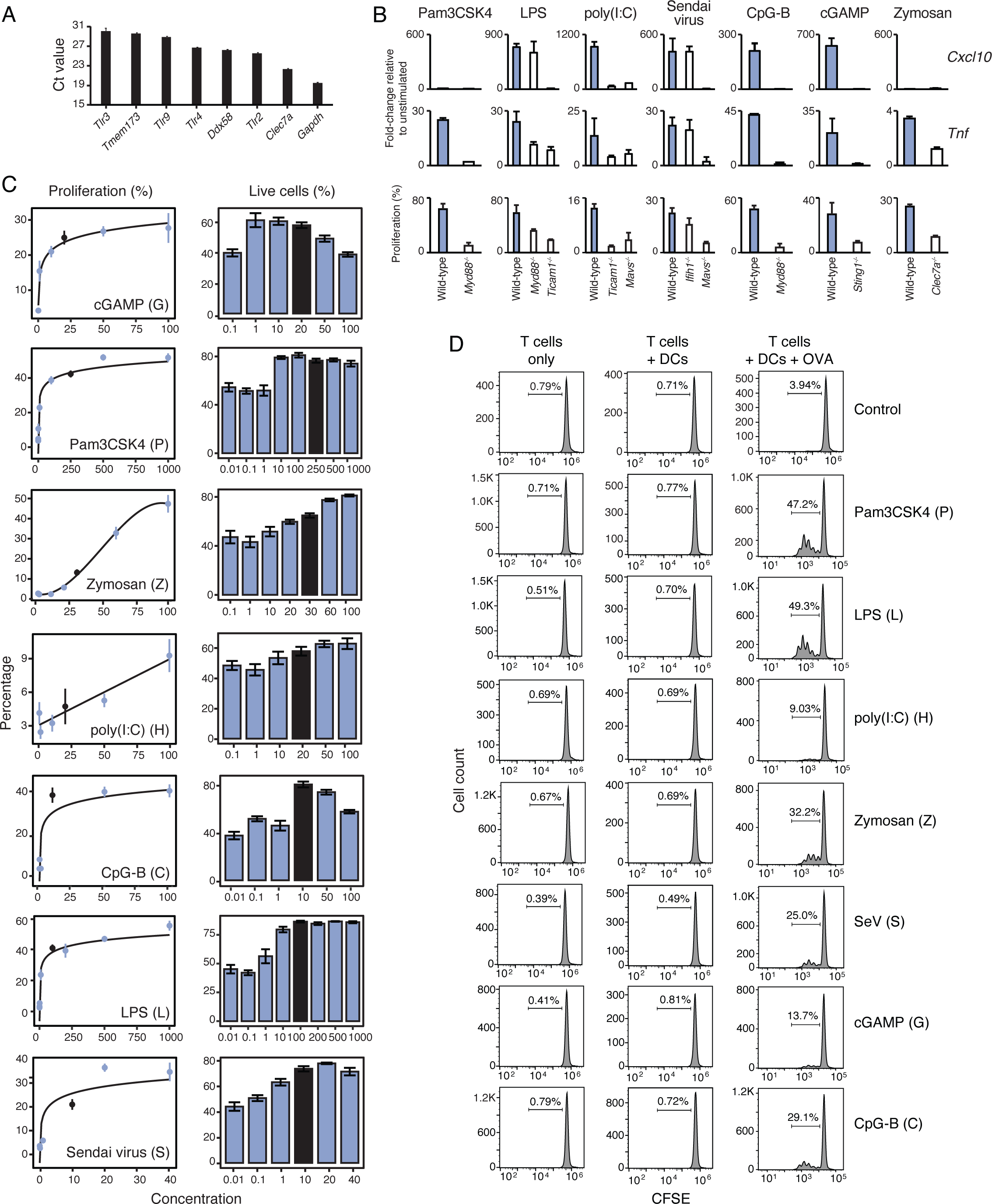
Selection of pattern-recognition receptor ligands. (A) Bar graph showing steady state mRNA expression in mouse dendritic cells (DCs) of the pattern recognition receptors or adaptor encoded by the following genes: *Tlr4*, *Tlr2*, *Tlr3*, *Tlr9*, *Ddx58* (RIG-I), *Clec7a* (DECTIN-1) and *Tmem173* (STING), which collectively recognize the seven ligands used in this study. Error bars, SD (*n* = 2). (B) Bar graphs showing mRNA expression of *Cxcl10* and *Tnf* (top two rows) and OT-II T cell proliferation (bottom row) upon stimulation of wild type (WT) and indicated knockout mouse dendritic cells (DCs) with the seven ligands selected for this study: cGAMP, CpG-B, Pam3CSK4, LPS, Zymosan, Sendai virus and poly(I:C). Error bars, SD (*n* = 2). (C) Dose-response analysis for the seven ligands selected for this study (indicated in left plots). For each ligand, indicated concentrations were used to stimulate DCs prior to adding OT-II cells (µg/mL for ligands G, Z, H, C and S; ng/mL for ligands P and L). Shown are proliferation (line plots; left) and viability (bar plots; right) measurements for OT-II T cells as a function of ligand concentrations. For proliferation plots, the logarithmic (G, C, P, L and S), linear (H) and cubic (Z) fits are shown as solid lines. The black circles (proliferation plots) and bars (viability plots) indicate the concentration selected for combinatorial screening analyses. Error bars, SEM (*n* = 3). (D) To control for the potential effects of ligands on T cells, as opposed to DCs, we measured OT-II T cells proliferation with CFSE in three conditions using indicated ligands: (1) T cells incubated with ligands only (T cells only; left), (2) T cells incubated with DCs that were stimulated and washed prior to the addition of T cells (T cells + DCs; middle), and (3) T cells incubated with DCs that were stimulated, pulsed with the ovalbumin protein and washed prior to the addition of T cells (T cells + DCs + Ovalbumin; right). Shown are representative CFSE profiles from live OT-II CD4^+^ T cells (*n* = 3).

**Figure S2.**
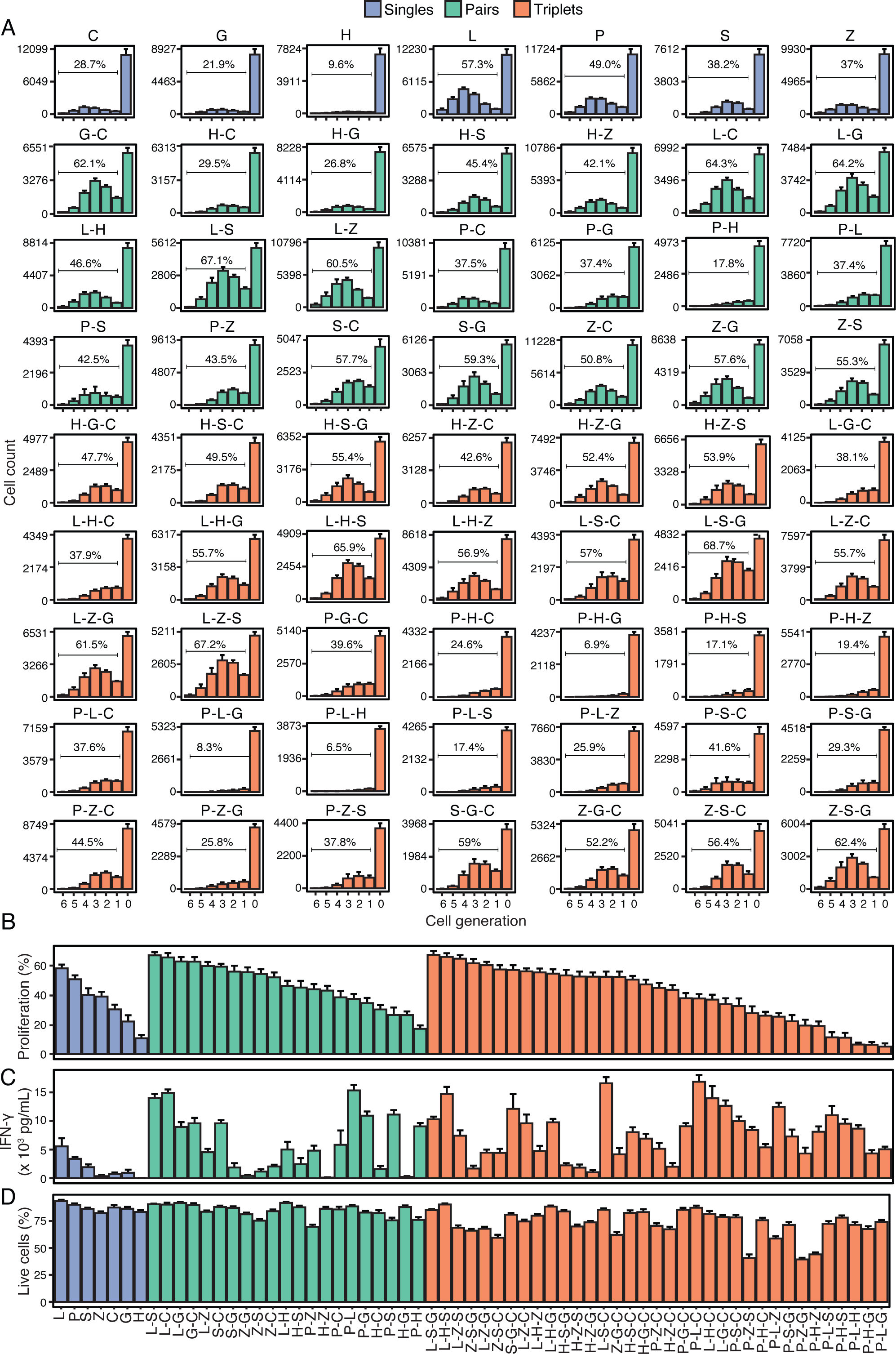
Impact of combinatorial stimulations of dendritic cells on T cell proliferation, cytokine secretion and viability. (A) Bar plots showing the CFSE profiles of OT-II cells cocultured with DCs stimulated with indicated ligand singles (blue), pairs (green) and triplets (orange). Percentages show the proportion of cells that underwent division. Error bars, SEM (*n* = 8). (B-D) Bar plots showing the proportion of OT-II cells that divided (B), the production of IFN-γ in DC-OT-II coculture supernatants (C), and the percentage of live OT-II cells (D) upon DC stimulations with indicated ligand singles (blue), pairs (green) and triplets (orange). Error bars, SEM (*n* = 8).

**Figure S3.**
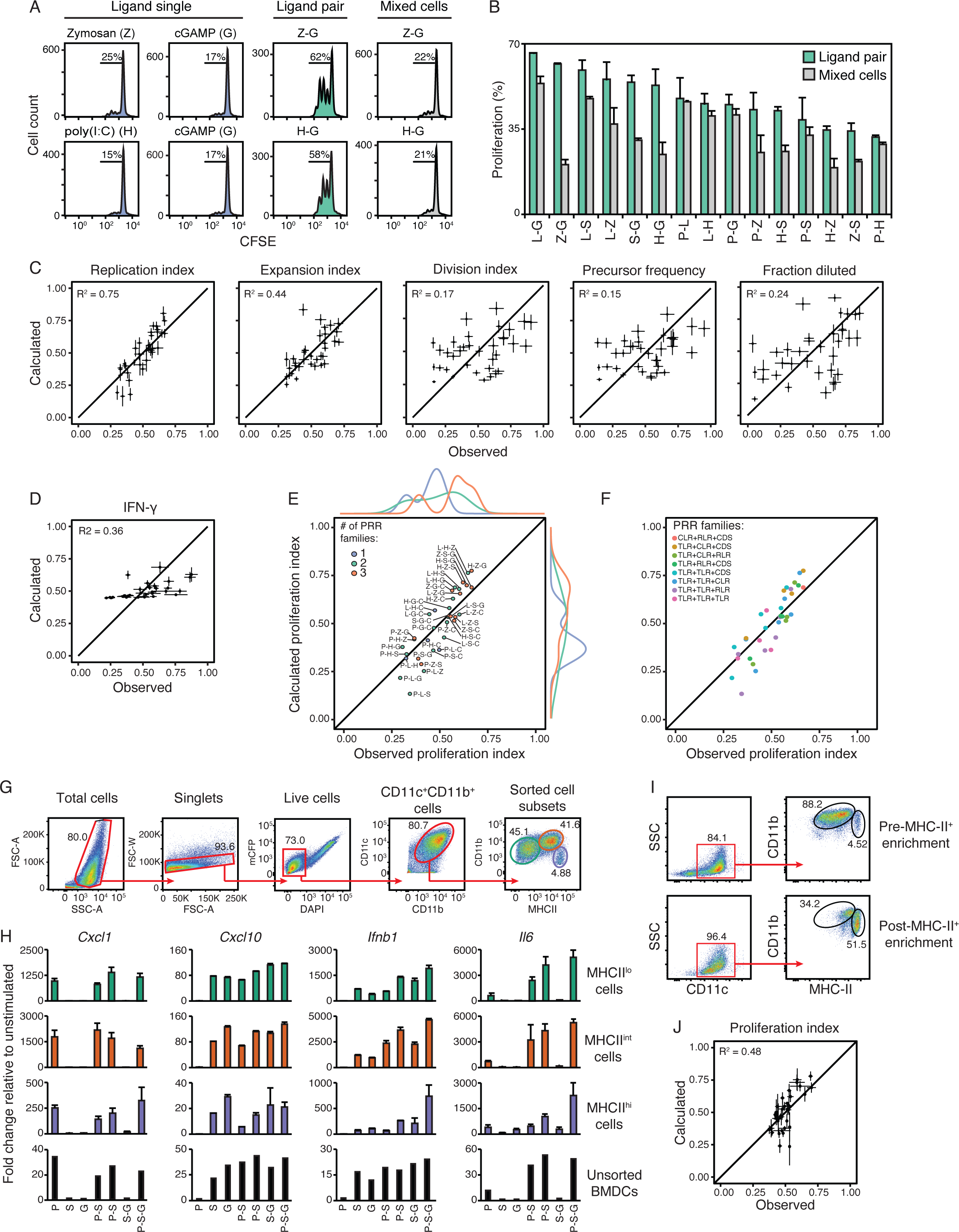
Assessing various T cell proliferation metrics and the clustering of ligand triplets and their proliferation in relationship to receptor families. (A) CFSE profiles of OT-II CD4^+^ T cells cocultured with DCs stimulated with indicated ligand singles, ligand pairs, or DCs stimulated with ligand singles and mixed at 1:1 ratio (Mixed cells). Percentages show the proportion of cells that underwent division. (B) Bar plots showing the proportion of OT-II cells that divided in cocultures with DCs stimulated with indicated ligand pairs or DCs mixed after single ligand stimulations (Mixed cells). Error bars, SD (*n* = 3). (C-D) Dot plots of the observed (X axis) and calculated (Y axis) triplet values for all 35 ligand triplets tested and using the indicated CFSE-derived proliferation metrics (see STAR Methods) (C), or IFN-γ concentrations from DC-OT-II cell cocultures (D). The solid line indicates y = x. Error bars, SEM (*n* = 8). (E-F) Scatter plots of the observed (X axis) and calculated (Y axis) triplet proliferation index values for all 35 ligand triplets tested (same data as shown in Figure 3B). The solid lines indicate y = x. Colors indicate the number (E) and the type (F) of PRR families covered by a given triplet. CLR, C-type lectin receptor; TLR, Toll-like receptors; RLR, RIG-I-like receptor; CDS, cytosolic dsDNA sensor. In D, density distributions are shown (top and right) for the number of PRR pathways targeted by a ligand triplet: one, two and three PRR families per triplet. (G-H) Gating strategy for fluorescence activated cell sorting (FACS) of the three following subpopulations present in GM-CSF-induced bone-marrow-derived DC cultures: CD11b^+^MHCII^lo^ (green), CD11b^+^MHCII^int^ (red), CD11b^+^MHCII^hi^ (purple) cells (G), and qPCR analysis of *Cxcl1*, *Cxcl10*, *Ifnb1* and *Il6* gene expression in indicated subpopulations (right) stimulated with indicated ligand singles or combinations (P, Pam3CSK4; S, Sendai virus; G, cGAMP) (H). The bottom row in G indicates changes in gene expression measured by RNA-seq on total, unsorted BMDCs. (I) Flow cytometry analysis of cells from total BMDC cultures before (top) or after (bottom) enrichment of MHC-II^+^ cells using magnetic microbeads. (I) Dot plots of the observed (X axis) and calculated (based on the Isserlis formula; Y axis) triplet proliferation index values for all 35 ligand triplets tested using MHC-II^+^ DCs enriched as shown in I. The solid line indicates y = x. Error bars, SEM (*n* =4-7).

**Figure S4.**
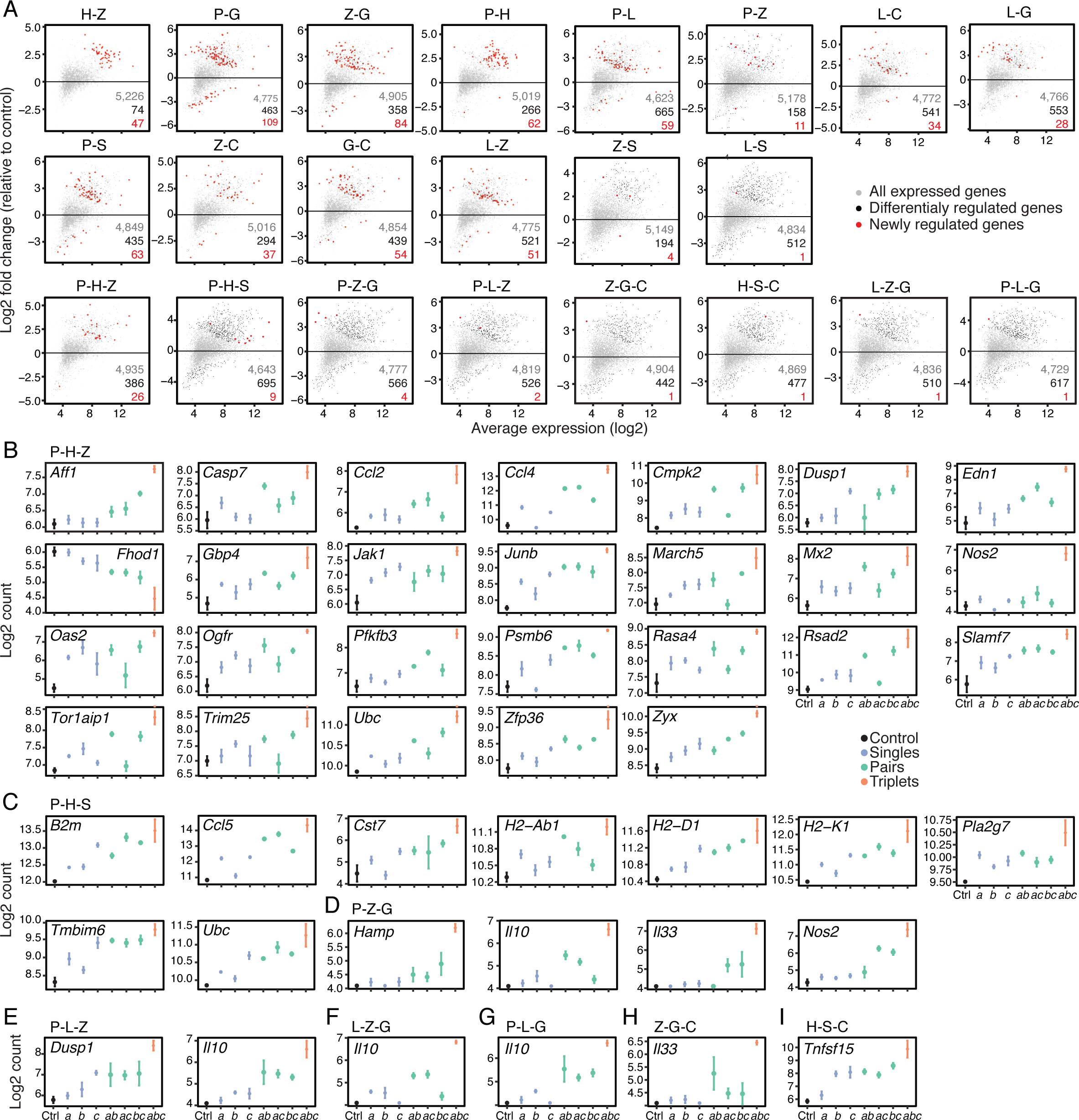
Genes newly regulated by ligand pairs and triplets compared to their composite ligand singles and pairs. (A) Dot plots showing log2 fold changes in gene expression (Y axis) in mouse DCs stimulated with indicated ligand pairs (14 out of 21 tested; top two rows) or triplets (8 out of 35 tested; bottom row) relative to unstimulated control cells against log2 average expression values (X axis) (*n* = 3). Red dots, genes regulated by indicated pairs or triplets but not by their composite ligand singles and/or pairs; black dots, all differentially regulated genes by indicated pairs or triplets; grey dots, all genes detected. (B-I) Shown are normalized expression levels in mouse DCs for indicated genes as log2 counts per million upon stimulation with indicated triplets: P-H-Z (B), P-H-S (C), P-Z-G (D), P-L-Z (E), L-Z-G (F), P-L-G (G), Z-G-C (H) and H-S-C (I). *a*, *b*, *c*, singles (blue dots); *ab*, *ac*, *bc*, pairs (green dots); *abc*, triplet (orange dots); Ctrl, control unstimulated cells (black dots). Error bars, SEM (*n* = 3).

**Figure S5.**
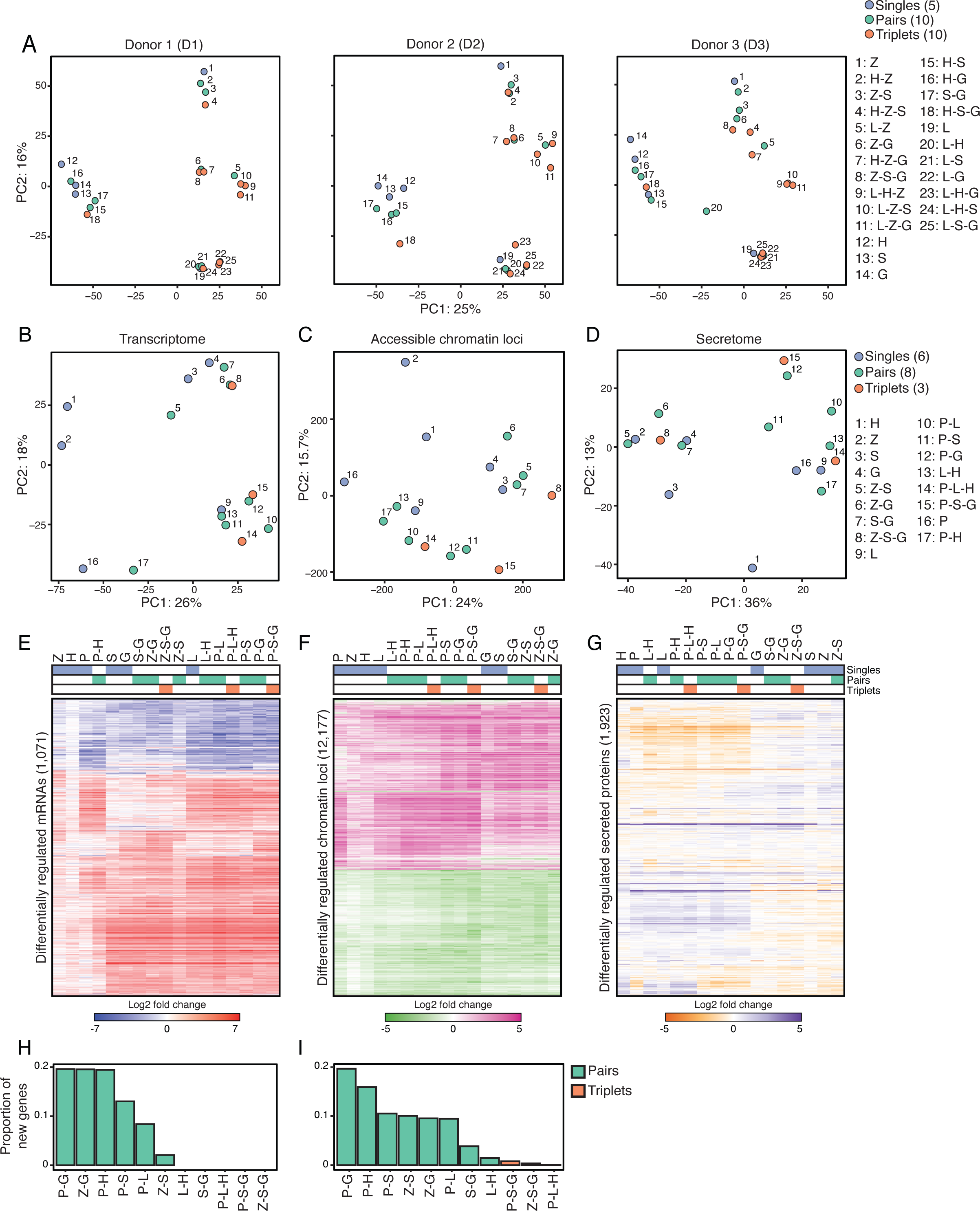
Effects of combinatorial stimulations on the transcriptional, chromatin accessibility and secretome profiles of dendritic cells. (A) Principal component analysis (PCA) of mRNA profiles of human blood monocyte-derived DCs (moDCs) from three independent healthy donors (from left to right: donor (D) 1 to 3) stimulated with indicated ligand singles (5; blue), pairs (10; green) or triplets (10; orange). (B-D) Principal component analysis (PCA) of mRNA (B), chromatin accessibility (C) and secretome (D) profiles of mouse DCs stimulated with indicated (numbers) ligand singles (6; blue), pairs (8; green) or triplets (3; orange) (*n* = 2-3). (E-G) Heatmaps of differentially regulated (rows) genes (E), accessible genomic loci (F), or secreted proteins (G) from mouse DCs stimulated with indicated ligand singles (6; blue), pairs (8; green) or triplets (3; orange). Values are log2 fold-changes relative to unstimulated cells (color scales) (*n* = 2-3). (H-I) Bar plots showing for each ligand pair (green) or triplet (orange) the proportion of genes (H) or accessible loci (I) regulated by a triplet or pair but not by its composite ligand singles and/or pairs relative to the total number of genes (H) or loci (I) regulated by that triplet (*n* = 2-3).

**Figure S6.**
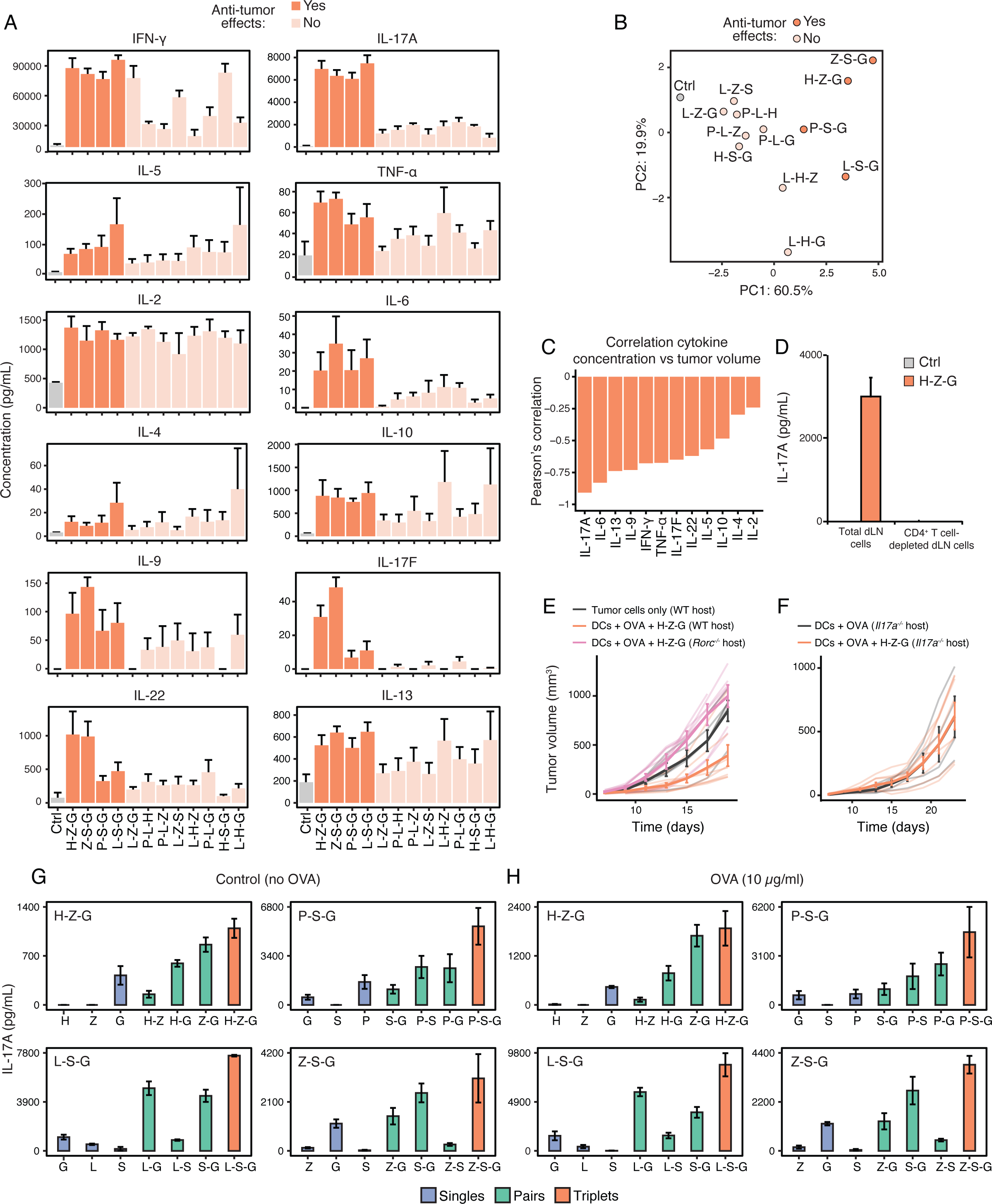
Effects of DC vaccines on draining lymph node cytokine secretion profiles. (A-B) Bar plots (A) and principal component analysis (PCA; B) of the secretion profiles of 12 cytokines measured in supernatants from total draining lymph node (dLN) cell cultures upon *in vitro* restimulation with the OVA protein (1 µg/mL). dLN cells were prepared from mice injected a week earlier with DC vaccines pulsed with OVA and stimulated with indicated ligand triplets or left unstimulated as control (grey). Dark and light orange bars (A) and dots (B) respectively indicate ligand triplets with or without anti-tumor properties *in vivo* (based on data shown in Figure 5). Error bars, SEM (*n* = 4). (C) Pearson’s correlation metric between cytokine concentrations (A) and B16-OVA tumor volumes at day 19 (Figure 5B) for the 12 adjuvant triplets used in tumor growth and dLN restimulation experiments. (D) Production of IL-17A by total dLN cell cultures prepared from mice injected a week earlier with DC vaccines pulsed with OVA and stimulated with the H-Z-G (poly(I:C)-Zymosan-cGAMP) triplet (H-Z-G) or left unstimulated as control (Ctrl). Shown are results for total dLN cells (total dLN) and dLN cells depleted of CD4+ T cells (CD4^+^ T cell-depleted dLN cells). (E-F) Mean tumor growth (solid lines) in cohorts of wild-type (WT), *Rorc^-/-^* and *Il17a^-/-^* knockout mice injected with 10^5^ B16-OVA and indicated DC vaccines. DCs + OVA, unstimulated DCs pulsed with OVA; DCs + OVA + H-Z-G, DCs stimulated with H-Z-G and pulsed with OVA. Light color lines indicate tumor growth for individual mice within each cohort. Error bars, SEM (*n* = 3-6 mice per cohort). (G-H) IL-17A production by total inguinal draining lymph node (dLN) cells from mice injected subcutaneously with DC vaccines loaded with OVA and stimulated with indicated ligand singles or combinations covering the following four triplets with anti-tumor effects: H-Z-G, L-S-G, P-S- G and Z-S-G. dLN cells were plated 7 days post-DC vaccination in medium without OVA (G), or restimulated with 10 µg/mL of purified OVA protein (H). Blue, singles; green, pairs; orange, triplets. Error bars, SEM (*n* = 4).

## SUPPLEMENTARY TABLES (separate Excel files)

Table S1. Data from cocultures of mouse DCs with transgenic CD4^+^ T cells

Average numbers of live transgenic CD4^+^ T cells (OT-II or SMARTA) in indicated CFSE peaks (0 to 6) and corresponding SEM values for all ligand singles, pairs and triplets (samples column) tested. Shown is data obtained by culturing total (A) or MHC-II^+^ (B) DCs with OT-II cells, and total DCs with SMARTA cells (C). IFN-γ concentration on coculture supernatants was measured by standard sandwich ELISA in A.

Table S2. mRNA expression data from mouse DCs stimulated with 7 singles, 21 pairs or 35 triplets of ligands for pattern recognition pathways

(A) Log2 fold-change values for the 1,357 genes (rows) differentially regulated in mouse DCs stimulated with ligand singles, pairs and triplets (columns) relative to unstimulated, control cells (FDR-adjusted p value < 0.01; *n* = 3).

(B) Binary table annotating the genes (rows) from A as newly regulated (1) by a ligand pair or triplet (columns) but not by matching composite ligand singles and/or pairs. Zeros indicate genes which are not newly regulated in a given pair or triplet.

Table S3. mRNA expression data from human DCs stimulated with 5 singles, 10 pairs or 10 triplets of ligands for pattern recognition pathways

(A) Log2 fold-change values for the 3,430 genes (rows) differentially regulated in human monocyte-derived DCs from three healthy donors (HD1-3) stimulated with ligand singles, pairs and triplets (columns) relative to unstimulated, control cells (FDR-adjusted p value < 0.01; *n* = 3).

(B) Binary table annotating the genes (rows) from A as newly regulated (1) by a ligand pair or triplet (columns) but not by matching composite ligand singles and/or pairs. Zeros indicate genes which are not newly regulated in a given pair or triplet.

Table S4. Accessible chromatin loci from mouse DCs stimulated with 6 singles, 8 pairs or 3 triplets of ligands for pattern recognition pathways

Log2 fold-change values for the 12,177 genomic loci differentially accessible (as measured by ATAC-seq) in mouse DCs stimulated with ligand singles, pairs and triplets (columns) relative to unstimulated, control cells (FDR-adjusted p value < 0.01; *n* = 2).

Table S5. Secretome data from mouse DCs stimulated with 6 singles, 8 pairs or 3 triplets of ligands for pattern recognition pathways

Log2 fold-change values for the 1,923 differentially regulated proteins from culture supernatants of mouse DCs stimulated with ligand singles, pairs and triplets (columns) relative to unstimulated, control cells (*n* = 2-3).

Table S6. Cytokine concentration measurements from draining lymph node (dLN) cell restimulation *in vitro* after DC vaccine injection subcutaneously

(A) Average concentration (pg/mL) and matching SEM (n = 4) values for indicated cytokines (12 in total) from the culture supernatants of total dLN cells harvested from mice injected with DC vaccines prepared with indicated ligand triplets (12 in total) or left untreated and pulsed with OVA only as control.

(B) Average concentration (pg/mL) and matching SEM (n = 4) values for indicated IL-17A from the culture supernatants of total dLN cells harvested from mice injected with DC vaccines prepared with indicated ligand singles, pairs or triplets.

